# Slope-Hunter: A robust method for index-event bias correction in genome-wide association studies of subsequent traits

**DOI:** 10.1101/2020.01.31.928077

**Authors:** Osama Mahmoud, George Davey Smith, Frank Dudbridge, Marcus Munafo, Kate Tilling

## Abstract

**Background:** Studying genetic associations with prognosis (e.g. survival, disability, subsequent disease events) is problematic due to selection bias - also termed index event bias or collider bias - whereby selection on disease status can induce associations between causes of incidence with prognosis. A current method for adjusting genetic associations for this bias assumes there is no genetic correlation between incidence and prognosis, which may not be a plausible assumption.

**Methods:** We propose an alternative, the ‘Slope-Hunter’ approach, which is unbiased even when there is genetic correlation between incidence and prognosis. Our approach has two stages. First, we use cluster-based techniques to identify: variants affecting neither incidence nor prognosis (these should not suffer bias and only a random sub-sample of them are retained in the analysis); variants affecting prognosis only (excluded from the analysis). Second, we fit a cluster-based model to identify the class of variants only affecting incidence, and use this class to estimate the adjustment factor.

**Results:** Simulation studies showed that the Slope-Hunter method reduces type-1 error by between 49%-85%, increases power by 1%-36%, reduces bias by 17%-47% compared to other methods in the presence of genetic correlation and performs as well as previous methods when there is no genetic correlation. Slope-Hunter and the previous methods perform less well as the proportion of variation in incidence explained by genetic variants affecting only incidence decreases.

**Conclusions:** The key assumption of Slope-Hunter is that the contribution of the set of genetic variants affecting incidence only to the heritability of incidence is at least as large as the contribution of those affecting both incidence and prognosis. When this assumption holds, our approach is unbiased in the presence of genetic correlation between incidence and progression, and performs no worse than alternative approaches even when there is no correlation. Bias-adjusting methods should be used to carry out causal analyses when conditioning on incidence.

## 1 Background

There is increasing interest in the use of genome wide association studies (GWAS) not only to investigate risk of disease, but to examine prognosis or outcome of people with the disease [l, 2, 3]. Studies of prognosis, of necessity, can be conducted only in those who have the disease, i.e. conditioning on disease incidence. This leads to a type of selection bias - termed index event bias or collider bias - whereby uncorrelated causes of the disease appear correlated when studying only cases [l, 2, 4]. This means that if there is unmeasured confounding between incidence and prognosis, then any cause of incidence will appear also to cause prognosis. Any cause of both incidence and prognosis will have a biased estimate of its effect on prognosis.

Figures l and 2 illustrate that a single nucleotide polymorphism (SNP), *G*, causing disease trait, *I*, becomes correlated with the confounder, *U*, of disease and subsequent trait, *P*, when conditioning on *I*. This induces association between *G* and *P* via the path *G* – *U* → *P* leading to index event bias in the SNP-prognosis association, if the confounding effects are not accounted for. If all causes of incidence were known and could be measured, the selection bias could then be removed, e.g. by using the inverse probability weighting (IPW) approach [5]. But, for IPW to be valid, the weighting model must be correctly specified, and must include all variables that are related to both incidence and to the variables in the analysis model (e.g. to prognosis and every genetic variant). However in most studies, these variables are not all known, and not all are measured.

The implications of index event bias have been addressed in several GWAS and MR studies [l, 2]. An example is the ‘paradox of glucose-6-phosphate dehydrogenase (G6PD) deficiency’ whereby among individuals selected according to their status of severe malarial anaemia (SMA), higher levels of G6PD deficiency appear to protect against cerebral malaria (CM) [6, 7]. A possible explanation is that if an individual with SMA has a high level of G6PD deficiency, they may well have lower levels of other risk factors for SMA. If lower levels of those other factors tend to decrease risk of CM, then the G6PD deficiency may appear to be protective against CM. In the notation of Figure 1, G6PD deficiency plays the role of the SNP *G*_1_ whereas *I* and *P* represent SMA and CM respectively. It has been suggested that this apparent protective effect is at least partially due to index event bias (collider bias) [8].

**Figure 1:**
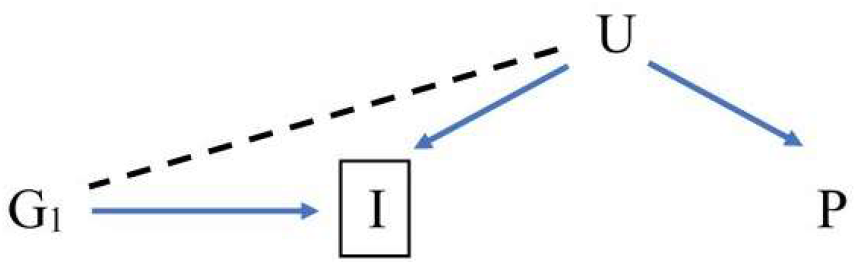
Directed acyclic graph for association of a SNP *G*_1_ with a prognosis trait *P* conditional on an incidence trait *I. U* is a composite variable including all common causes of *I* and *P*, including polygenic effects as well as non-genetic factors. Conditioning on *I* induces the association between *G*_1_ and *U*, shown by the dashed line, leading to biased association between *G*_1_ and *P* via the path *G*_1_ − *U* → *P*. The association of *G*_1_ with the prognosis *P* when conditioning on incidence is entirely due to the index event bias

A method for adjusting genetic associations for the index event bias has been proposed whereby estimated residuals from the regression of SNP-prognosis associations on SNP-incidence associations give bias-adjusted effects on prognosis [4]. This method assumed that the direct genetic effects on incidence and prognosis are linearly uncorrelated. But, this assumption may be incompatible with the premise of genetic studies in which shared pathways of incidence and prognosis have been observed for many traits. For example, such shared pathways may be common for psychiatric traits [9], metabolites [10] and phenotypes related to cumulative effects of long-term exposures [11].

We propose a novel method, referred to as ‘Slope-Hunter’, for adjustment of index event bias in GWAS of progression studies with potentially correlated direct genetic effects on incidence and prognosis. This is achieved by first identifying the class of SNPs which only affect incidence, and using these to obtain an unbiased estimate of the correction factor that is then used to adjust for the bias for all genetic variants. We evaluate the Slope-Hunter method by comparing its type-1 error, power and bias with the previously proposed methods in an extensive simulation study with realistic parameters.

## 2 Methods

For an individual SNP, it is assumed that a continuous incidence trait *I* is linear in the coded genotype *G*, common causes *U* of incidence and prognosis, and unique causes *ε*_*I*_ of *I*:

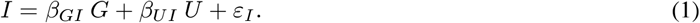

Moreover, we assume that a continuous prognosis trait *P* is linear in *G, U*, and unique causes *ε*_*P*_ of *P*, with an additional linear effect of *I*:

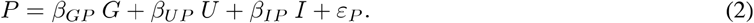

The effect of our interest is the SNP effect on prognosis, *β*_*GP*_, conditional on incidence *I* and confounders *U.* However in practice, we can only estimate the SNP-prognosis association conditional on incidence, denoted by 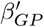 in equation 3, because all relevant confounders may not be observed.

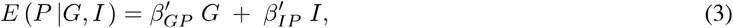

where 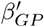 is a biased estimate of SNP effect on prognosis (termed ‘conditional estimate’), whereas 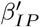 is a biased estimate of the causal effect of incidence on prognosis. Equations 1 and 2 assumed that both *I* and *P* are continuous traits. However, if *I* and/or *P* are binary, as in the case for disease traits, it has been shown that the logistic and probit link functions are approximately linear for small effects, as typically is the case for polygenic traits [4]. Therefore, we still consider the linear models presented in equations 1 and 2. If the incidence is a binary disease trait, then conditioning on the incidence, as in equation 3, is equivalent to analysing prognosis among cases only:

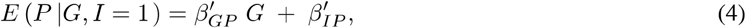

Dudbridge et. al. (2019) showed that the conditional estimate 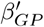 can be formulated as the true effect *β*_*GP*_ plus a bias that is linear in the SNP effect on incidence *β*_*GI*_ [4]. In particular, the conditional estimate can be expressed as follows:

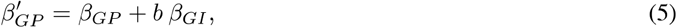

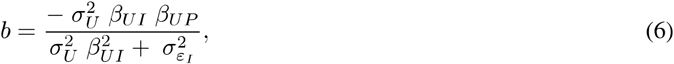

where 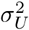 and 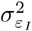 are variances of confounders and residual unique causes of *I* respectively. Therefore, by regressing the conditional estimates 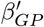 on *β*_*GI*_ for all SNPs, the slope, *b*, could be estimated using ordinary least squares (OLS), assuming that [4]:

- *A*_1_: The effects of SNPs on incidence are linearly uncorrelated with their direct effects on prognosis (i.e. index coefficient linearly uncorrelated with direct effect, referred to as the ‘InCLUDE’ assumption).
- *A*_2_: The confounder effects - and hence *b* - are constant across all SNPs.

The estimated slope, 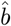, can then be used to obtain bias-adjusted effects on prognosis for each SNP by calculating the residuals of equation 5 as follows:

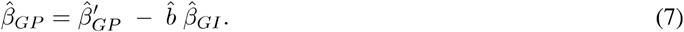

If there are shared pathways for both incidence and prognosis whereby the direct effects on prognosis are correlated with effects on incidence, e.g. as shown in Figure 3, then the assumption *A*_1_ can be violated producing bias in 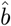, and hence not correcting adequately for the index event bias, see Figure 4. Although the non-genetic component of *U* is constant across all SNPs by definition, assuming it has no interaction with each SNP, the genetic component of *U* may differ across SNPs leading to violation of assumption *A*_2_. For a single SNP *G*_1_ affecting incidence only (Figure 1), the genetic component of *U* is the entire shared genetic basis of *I* and *P.* Whilst for a single SNP *G*_2_ affecting both incidence and prognosis (Figure 2 and Figure 3, the genetic component of *U* equals entire shared genetic basis of *I* and *P*, minus the component attributed to the SNP under consideration, *G*_2_.

In this paper, we propose a novel approach for adjustment of index event bias in GWAS of progression studies, referred to as ‘Slope-Hunter’. It firstly identifies variants affecting neither incidence nor prognosis, i.e. the class of *G*_4_ SNPs as shown in Figure 5(b); and variants affecting prognosis only, i.e. the class of *G*_3_ SNPs as shown in Figure 5(a). These classes of variants do not have an effect on incidence, and therefore do not suffer bias. A random sub-sample of *G* _4_ SNPs is retained in the analysis to facilitate pattern identification, and the remaining SNPs of these groups are excluded. Then, the pattern of the class of variants affecting incidence only, *G*_1_, is distinguished and used to obtain an unbiased estimate of the slope *b*.

**Figure 2:**
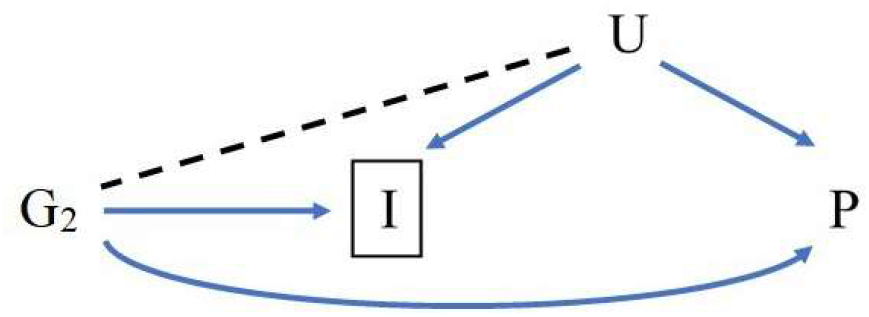
Directed acyclic graph for association of a SNP *G*_2_ with a prognosis trait *P* conditional on an incidence trait *I. U* is a composite variable including all common causes of *I* and *P*, including polygenic effects as well as non-genetic factors. Conditioning on *I* induces the association between *G*_2_ and *U*, shown by the dashed line, leading to association between *G*_2_ and *P* via the path *G*_2_ − *U* → *P*, in addition to the direct effect *G*_2_ → *P*.

**Figure 3:**
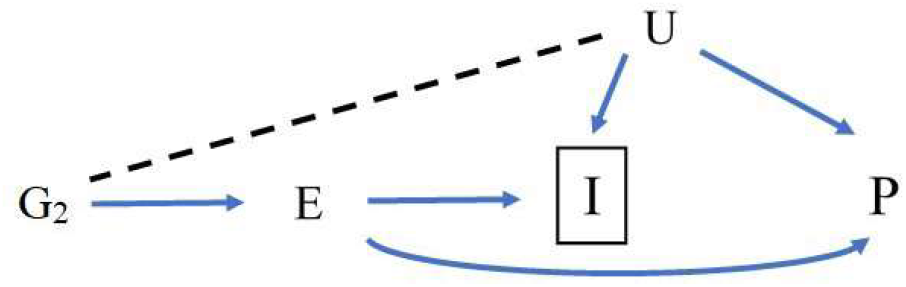
Directed acyclic graph for association of a SNP *G*_2_ with a prognosis trait *P* conditional on an incidence trait *I. U* is a composite variable including all common causes of *I* and *P*, including polygenic effects as well as non-genetic factors. There is a shared pathway for both incidence and prognosis via an exposure *E* leading to correlation between effects on prognosis and effects on incidence. Conditioning on *I* induces the association between *G*_2_ and *U*, shown by the dashed line, leading to association between *G*_2_ and *P* via the path *G*_2_ − *U* → *P*, in addition to the effect *G*_2_ − *E* → *P*.

**Figure 4:**
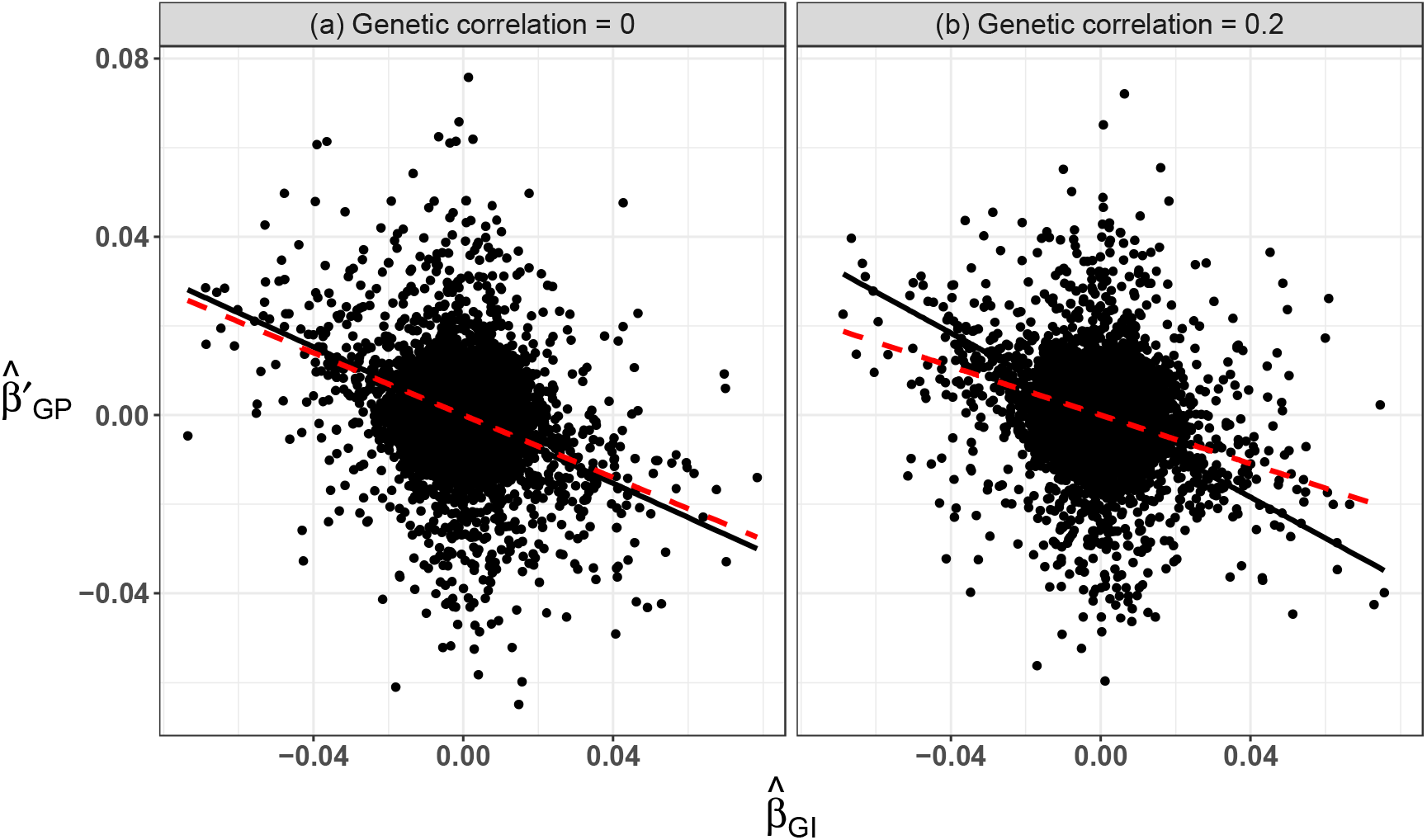
Scatter plots for estimates of SNP-incidence associations, 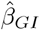, and conditional estimates, 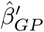, simulated from dataset of 20,000 ind i v iduals for l 0,000 independent SNPs, with: (a) no genet ic correlation betwee n SNP effec ts on incidence and prognosis; (b) correlated genetic effects (correlation coefficient = 0.2). Five-percent of SNPs have effec ts on incidence only, 5% on prognosis only, and 5% on both. Heritability of incidence and prognosis is 50% and non-genetic common factors of explain 40% of variation in both incide nc e and prognosis. These simulations induced index even t bias due to confounders that explain 40% and 60% of variation in prognosis in (a) and (b) respectively. The true confounding effec ts are represented by slopes of the black solid lines, (a) −0.38 and (b) −0.46, while the esti mated con ection factors using Dudbridge et. al. (2019) meth od are represented by slopes of the red dashed lines, (a)-0.35 and (b)-0.27. This figure illus trates potential inadequate conection usi ng the Dudbridge et. al. (2019) method when the ‘InCLUDE’ assumption (Index Coefficient Linearly Unconelated with Direct Effect) is violated.

**Figure 5:**
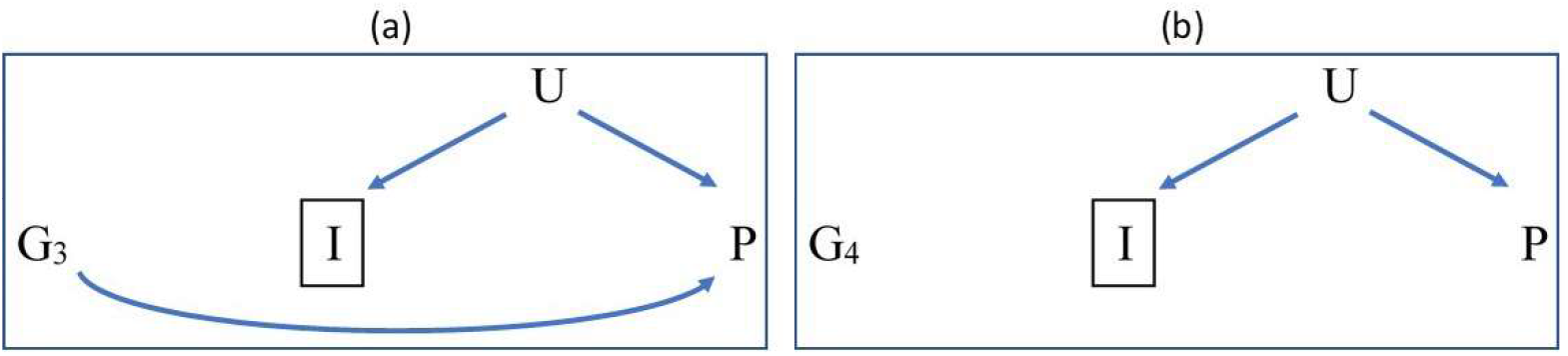
Directed acyclic graph for association of: (a) a SNP *G*_3_ with a direct effect on a prognosis trait *P*, with no effect on incidence *I;* (b) a SNP *G*_4_ with effect on neither *I* nor *P*, conditional on an incidence trait *I. U* is a composite variable including all common causes of *I* and *P*, including polygenic effects as well as non-genetic factors. Conditioning on *I* does not induce biased association between either *G*_3_ or *G*_4_ and *P* as no association between *G*_3_ or *G*_4_ and *U* are produced since either SNP does not affect *I*.

### The ‘Slope-Hunter’ Method

Equation 5 can be reformulated according to SNP associations with incidence and prognosis as follows:

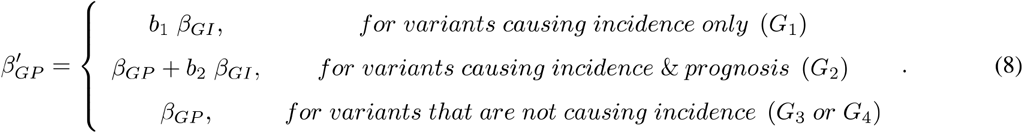

For the class of variants only affecting incidence, *G* _1_, assumption *A*_1_ is not violated, since these SNPs have no direct effects on prognosis. In addition, the genetic and non-genetic components of *U*, and hence the slope *b*_1_ is constant across all elements of the class *G*_1,_ i.e. assumption *A*_2_ is satisfied. Therefore, regressing the conditional estimates of SNP-prognosis associations, 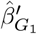 on estimates of SNP-incidence associations, 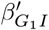, enables us to obtain an unbiased estimate,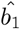, for *b*_1_, see equation 8, that can be utilised as an adjustment factor. Although the unbiased estimate of this adjustment factor is obtained by using the class of variants only affecting incidence, *G*_1_, it can still be used to correct bias for all SNPs by substituting 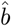 with, 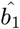 in equation 7, assuming that *b*_1_ ≈ *b*_2_, which is typically the case for small effects as in GWAS.

For *G*_2,_ either the assumption *A*_1_, *A*_2_ or both may not be satisfied, e.g. if there are underlying shared biological pathways for *I* and *P* and/or there are major variants accounting for substantial covariation in *I* and *P* leading to non-constant genetic component of *U.* In such cases, the slope estimated using the *G*_2_ class of variants, 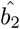 might be biased. Figure 2 shows the DAG for a SNP *G*_2_ that has a direct effect on both incidence and prognosis, whereby the estimated SNP-prognosis association conditional on incidence, 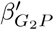 is due to the collider bias in addition to the SNP’s direct effect on prognosis. In such a case, the ratios of 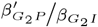 are not constant across all elements of the set of *G*_2_ SNPs.

For *G*_3_ and *G*_4_, the estimated direct effects on prognosis do not suffer from the index event bias, since conditioning on *I* does not induce association between *U* and *G* unless the genetic variant *G* causes the incidence, see Figure 5.

The Slope-Hunter method utilises cluster-based models [12, 13, 14] to identify the class of *G*_1_ and then adjust for the index event bias. The pseudo code of the ‘Slope-Hunter’ approach is presented in Algorithm l and a graphical illustration of the phases of our approach is presented in Figure 6.

**Figure 6:**
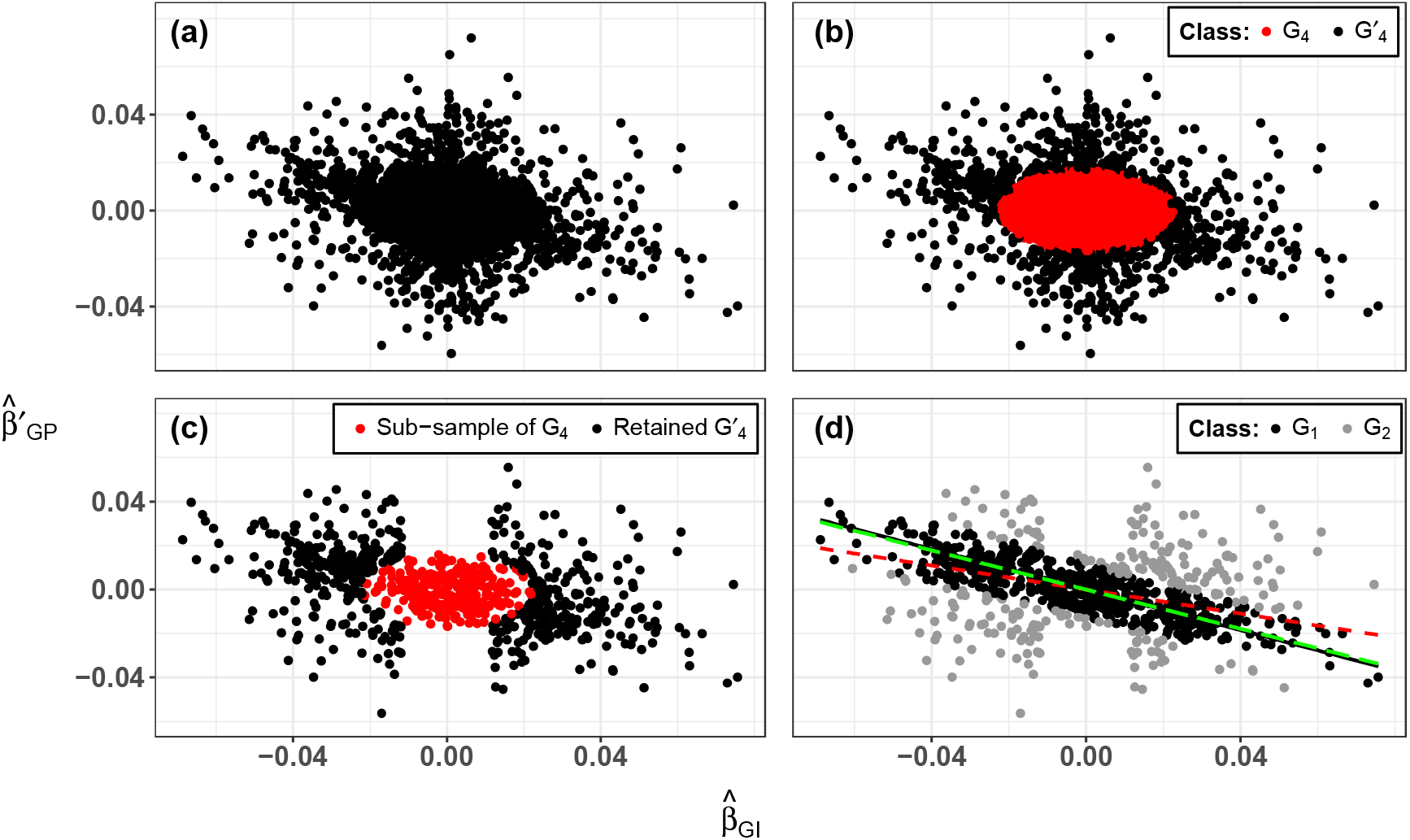
A graphical illustration for phases of the Slope-Hunter approach: (a) inputs of SNP-incidence associations, 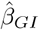, and conditional estimates, 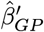 shown in Figure 4(b), that were si mulated with correlated genetic effec ts (coITelation coefficient= 0.2) on incidence and prognosis; (b) Slope-Hunter identifies variants affecting neither incidence nor prognosis, i.e. the class of *G*_4_ SNPs; (c) a random sub-sample of *G*_4_ SNPs is retained in the analysis, whereas the remaining variants in this group and the variants affecting prognosis only, i.e. the class of *G*_3_ SNPs, are excluded. The latter is identified using a *p*-value threshold for SNP-incidence associations; (d) The class of variants affec ting incidence only, *G*_1_, is identified and an estimate of its linear regression slope (represented by green long-dashed line, slope = −0.445) is obtained to cmTect for the index event bias of all SNPs. The true confounding effect is represented by the black solid line’s slope, −0.460, whereas the estimat ed coITection factor usi ng Dudbridge et. al. (2019) method is represented by the red dashed line’s slope, −0.273.

### Identification of variants affecting incidence only

Our proposal requires summary-level GWAS statistics and their standard errors for an incidence trait 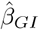 and *s*_*GI*_; and a prognosis trait *P* conditional on 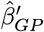 and 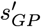. These inputs are obtained from GWAS of incidence and prognosis conditional on incidence. Other inputs provided by the user are an ordered set Λ of *k* proportions representing sub-sample sizes *λ*_*j*_, *j* = 1,…, *k* of the class *G*_4_ that are retained in the analysis; a *p*-value threshold *η* for identifying SNPs with no effect on incidence, the class *G*_3_; and a tolerance distance scalar *δ* bused for validating the cluster solutions. Our developed algorithm uses default values for these parameters, Λ = {3%, 4%, …, 10%}, *η* = 0.1 and *δ* = 1, that performed the best in our simulation studies. The Slope-Hunter approach produces the bias-adjusted estimates of SNP-prognosis associations 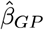 and their estimated standard errors *s*_*GP*_.

Since the adjustment factor, *b*_1_, should be estimated using a set of independent SNPs, GWAS are first pruned by linkage disequilibrium (LD). The first step of our approach, presented in Algorithm l, uses the pruned GWAS statistics for the incidence *I* to obtain *p*-values for SNP-incidence associations, *P*_*GI*_ (line 1). Then, we fit a bivariate normal mixture model with two components to estimate probabilities, and to assign memberships, of the cluster *G*_4_. Each mixture component is specified as a two-dimensional ellipse with varying geometric features: area, denoted by *ζ* (i.e. 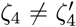); shape, denoted by 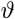 (i.e. 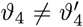); orientation, denoted by *θ* (i.e. 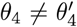), between the class *G*_4_ and its complementary cluster *G*′_4_ (line 2). The geometric features of each cluster are determined by its covariance matrix that can be decomposed to the form of 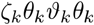 for a cluster *k*, where *ζ*_*k*_ is a scaler controlling the cluster area, 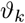 is a 2 × 2 diagonal matrix whose determinant equals 1 characterising its shape, and *θ*_*k*_ is a 2 × 2 orthogonal matrix characterising the orientation of corresponding ellipsoid [15, 16]. Figure 7 illustrates the geometric features of clusters identified by the Slope-Hunter method using the data simulated for Figure 4. Since SNPs of the class *G*_4_ have effects on neither incidence nor prognosis by definition, then their corresponding pairs of association coefficients 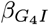 and 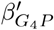 should have no structural function with respect to neither incidence nor prognosis. Hence their estimated values are expected to scatter around the origin in a noisy form with a high probability concentration, since the majority of GWAS variants are expected to belong to this class, see Figure 6(b). The *G*_4_ class is then identified as the cluster that contains the point with the smallest Euclidean distance from the origin.

**Figure 7:**
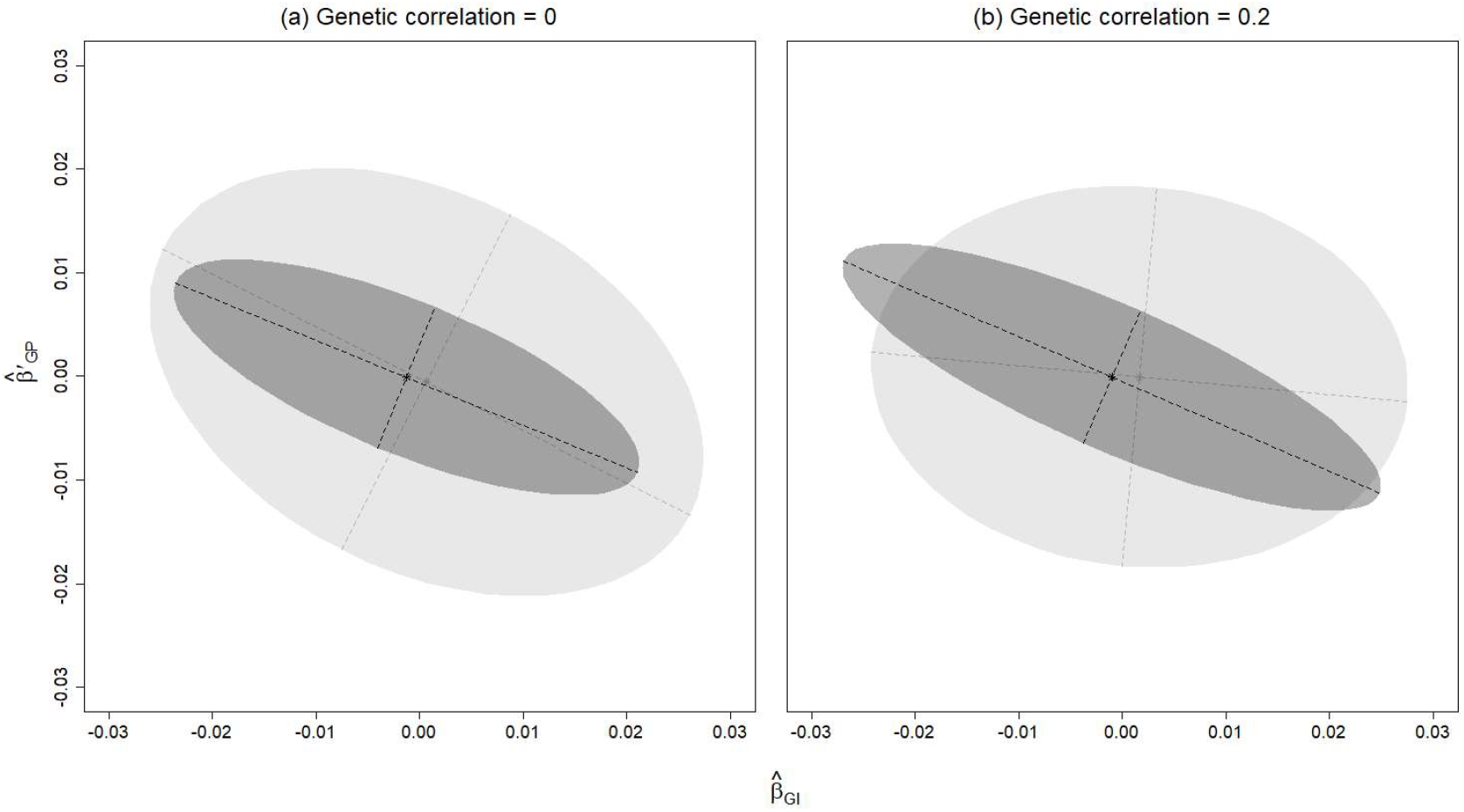
Illustration of mixture components (clusters) estimated by the Slope-Hunter method for the class of variants affecting incidence only (darker colour) and the calss of variants affecting both incidence and prognosis (lighter colour) based on values of SNP-incidence associations, 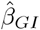, and conditional estimates, 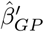 simulated from dataset of 20,000 individuals for 10,000 independent SNPs, with: (a) no genetic correlation between SNP effects on incidence and prognosis; (b) correlated genetic effects (correlation coefficient = 0.2). Each cluster is ellipsoidal with geometric features: area (coloured); shape (of the estimated ellipse); orientation (illustrated by orthogonal dashed lines, i.e. axes of the ellipse), that are determined by its covariance matrix [15, 16]

Exclusion of all SNPs assigned to the class *G*_4_ may discontinue the dimensional space leading to poor or invalid clustering of the remaining data. Therefore, we retain a fraction, *λ* of *G*_4_ SNPs to facilitate pattern recognition of the class *G*_1_. The selection of this sub-sample is performed randomly using a weighted score defined as a modified form of the Euclidean distance from the origin, *w*_*g*_, for each SNP *g* ∈ *G*_4_ (lines 3-5). This implies that data points with larger distance from the origin, with higher weights given to the prognosis dimension, are more likely to be retained in the analysis. The weights are then normalised to lie within [0, 1] (line 6).

We define C as a set of the adjustment factor candidates. This is initialised as an empty set (line 7) that will be iteratively updated by assigning a candidate adjustment factor 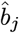 obtained by fitting a clustering solution *f*_*j*_ using a different sub-sample size *λ*_*j*_ ∈ Λ, at each iteration, *j* = 1, …, *k*. For each given proportion *λ*_*j*_, the following steps are performed:

1. A sub-sample of SNPs, *ℓ*_*j*_, of size *λ*_*j*_ is randomly selected from the class *G*_4_ using the vector of weights *ω*_*G*_4__ whose individual element for a single SNP *g* is *ω*_*g*_ (line 9).
2. The set of SNPs, *G\**, retained in the analysis is defined as the selected sample *ℓ*_*j*_ in addition to all SNPs *g* ∈ *G*′_4_ whose *p*-values *P*_*gI*_ < *η* (line 10). The latter procedure is designed to exclude SNPs that have no effects on *I* among the cluster *G*′_4_ (i.e. it excludes the *G*_3_ group).
3. SNPs of the *G*_1_ class should differ from the *G*_2_ class in terms of their patterns of estimated values around the true slope *b*_1_, because the *G*_1_ SNPs satisfy a proportional relationship between *β′*_*GP*_ and *β*_*GP*_, whilst SNPs of the *G*_2_ class deviate from such a proportional form with magnitudes dependent on true size of effect on prognosis for each SNP, *β*_*GP*_, see equation 8. The data points of *G*_1_ and *G*_2_ sets can then be treated as observations generated by two unknown distinct bivariate normal distributions. The Slope-Hunter method fits a cluster-based model (line 11) using the expectation-maximization (EM) algorithm to identify the underlying distributions and to estimate probability of each SNP belonging to the *G*_1_ class. For the set of SNPs *G\**, we fit a bivariate normal mixture model with two components, *G*_1_ and *G*_2_ that are ellipsoidal with varying orientations (*θ*_1_ ≠ *θ*_2_), representing varying slopes for *G*_1_ and *G*_2_, but possibly with equal areas and shapes. This is achieved by employing the model with the minimum Bayesian information criterion (BIC) among models with: equal areas and shapes 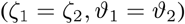; varying areas and equal shapes 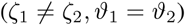; equal areas and varying shapes 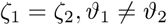; varying areas and shapes 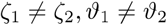.
4. We define 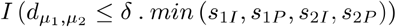 as an indicator that sets to 1 if 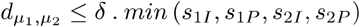, otherwise it sets to zero (line 12); where 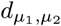 is the Euclidean distance between the means *μ*_1_ and *μ*_2_ of the identified clusters *G*_1_ and *G*_2_, the *s*_1*I*_, *s*_1*P*_, *s*_2*I*_, *s*_2*P*_ represent standard deviations of the clusters *G*_1_ and *G*_2_ respectively on each dimension *I* and *P*. The scalar, *δ* is set to scale the minimum standard deviation of the clusters. For example, if *δ* = 1 (the default), then the obtained cluster solution will be considered as a candidate for bias adjustment only if the distance between means of its components is not larger than the minimum standard deviation *s*_*ij*_, *i* = 1, 2, *j* = *I, P*, i.e. the value of parameter *δ* allows users to control how to trim the clustering solution candidates.
5. Since the class *G*_1_ is assumed to be scattered around its slope with less variation than the other class *G*_2_, then the Class *G*_1_ is identified as the one producing the minimum standard error of the slope *b*_*j*_ when regressing SNP-incidence on SNP-prognosis associations. If the model *f*_*j*_ produces a valid clustering candidate, as indicated by line 12, then a linear regression model *M*_*j*_ regressing associations with incidence, 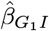 on association with prognosis 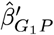 for SNPs in the class *G*_1_ is fitted (line 13).
6. The set 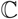 is then updated by adding the element representing the estimated slope 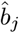 from the model *M*_*j*_ (line 14).

The optimal adjustment factor, 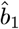, is then identified as the slope 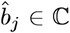 with the minimum standard error (line 17). Since analyses are conducted on finite sample estimates 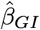 and 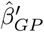 the regression may yield an estimate 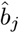 that is biased towards zero. Therefore, we have adjusted for such a regression dilution using the Hedges-Olkin estimator [4]. The bias-adjusted estimates of effects on prognosis and their standard errors for all SNPs *g* ∈ *G* can then be obtained using the optimal adjustment factor 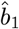 after regression dilution correction (lines 19, 20).

All procedures described in this manuscript have been implemented into an open source R package named ‘Slope-Hunter’ that would be available from https://github.com/Osmahmoud/Slope-Hunter

#### Algorithm 1 Slope-Hunter Adjustment for index event bias in GWAS of subsequent events

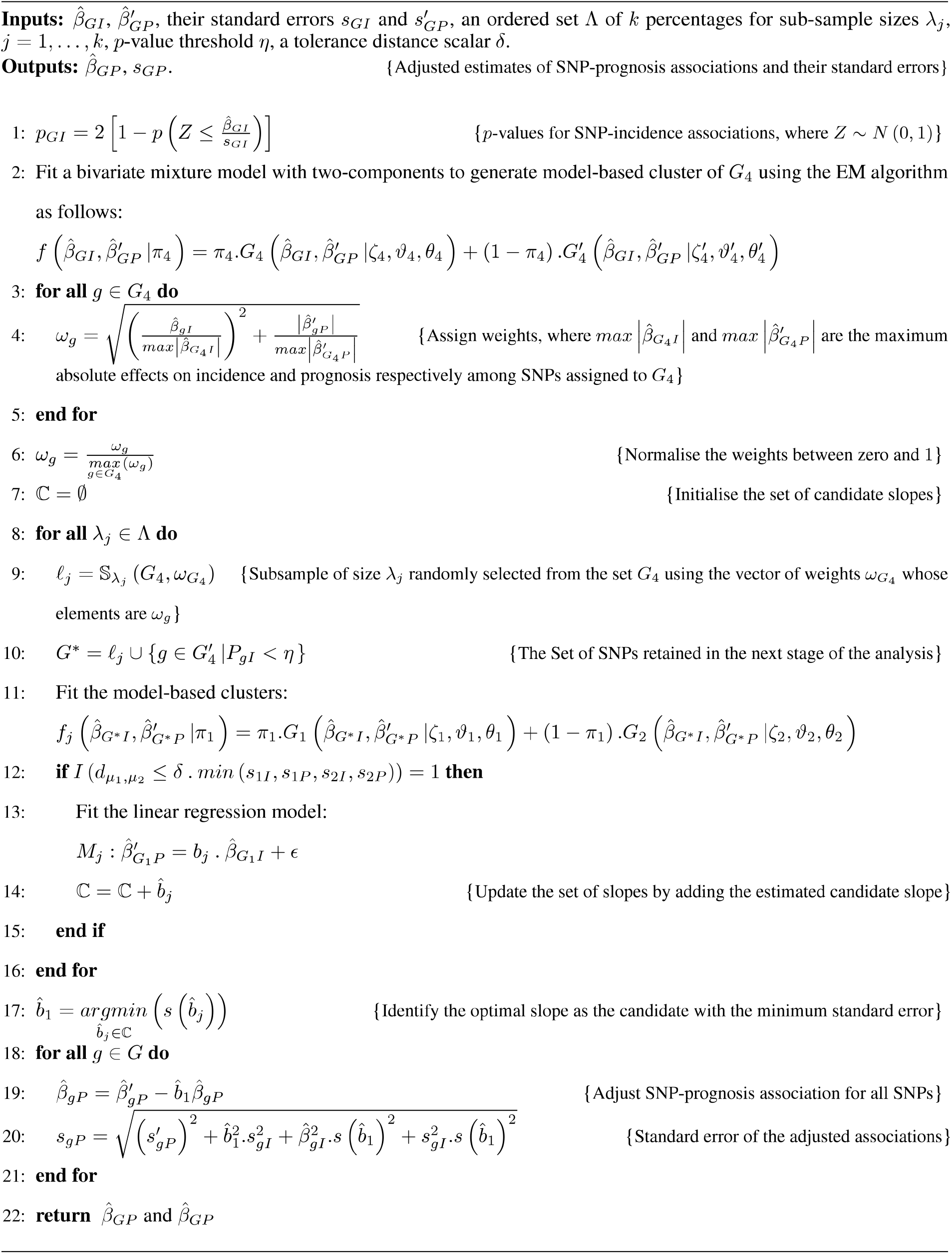

### Underlying assumptions

Our analytic approach assumes the analysed SNPs are independent, do not interact with the confounders, and have linear effects with the incidence and prognosis. Although our framework assumed a linear effect of *I* on *P*, equation 2, the size of that effect is not important for our theoretical developments and it might be zero.

The cluster-based model technique theoretically identifies a component distribution that is concentrated around a line (within two-dimensional settings as in our context), which is the first principal component of the corresponding cluster [12]. By definition, the SNP-prognosis associations for genetic variants of the class *G*_1_ are a function of their effects on incidence, see equation 8. Our procedure assumes that the variance of incidence explained by the class *G*_1_ is at least as much as that explained by the class *G*_2_. If this variance condition is true, then the data points of the true class *G*_1_ are more concentrated around the true slope *b*_1_ than the class *G*_2_. In this case, the pattern of the *G*_1_ class can be unambiguously identified by the cluster-based model, and the Slope-Hunter is theoretically valid. This assumption is a less restrictive case of the zero modal pleiotropy assumption (ZEMPA) [17] used in the MR context in which the number of valid instruments are assumed to be larger than any other group sizes of invalid instruments with unique estimates of the causal effect.

Since any correlation between incidence and prognosis should be included in *U* by definition [4], then our method is robust against the overlap between samples in prognosis GWAS and incidence GWAS. Thus, it is appropriate for one-sample analysis of both incidence and prognosis.

### Simulations

We simulated five scenarios, each with 10, 000 independent SNPs under Hardy-Weinberg equilibrium with minor allele frequencies drawn from a uniform distribution over the interval [0.01, 0.49]. For all scenarios, both incidence and prognosis were simulated as quantitative traits. The heritability under models shown in equations 1 and 2 was 50%, and the non-genetic confounder, *U*, explained 40 % of variation in both incidence and prognosis, with positive coefficients, *β*_*UI*_ and *β*_*UP*_ SNP effects, confounders and residual variation, *ε*_*I*_ and *ε*_*P*_, were drawn from normal distributions. Data were simulated for 20, 000 unrelated individuals.

In the first scenario (Sc.1), 1% of SNPs (100 SNPs) had effects on incidence only (*G*_1_) explaining 45 % of its variation, 9 % had effects on both incidence and prognosis (*G*_2_) explaining 5% of variation in incidence, and 1% had effects on prognosis only (*G*_3_). This set up of simulation implies that an index event bias is developed from confounders, with genetic and non-genetic components, that together explain 0.05 + 0.40 = 45% of variation in prognosis. In the second scenario (Sc.2), 5% of SNPs represented *G*_1_, 5% represented *G*_2_, each explaining 25% of variation in incidence, and 5% of SNPs represented *G*_3_. This simulation implies that an index event bias is developed from confounders that explain 0.25 + 0.40 = 65% of variation in prognosis. This simulation reflects a similar scenario to the one discussed by Dudbridge et al [4]. In the third scenario (Sc.3), 1 % of SNPs represented *G*_1,_ and 9% represented *G*_2,_ each explaining 25% of variation in incidence, whereas 1% of SNPs represented *G*_3_. This implies that index event bias is developed from confounders that explain 65% of variation in prognosis. The fourth scenario (Sc.4) had 3% of SNPs representing *G*_1_, 7% representing *G*_2_, explaining 15% and 35% of variation in incidence respectively, and 3% representing *G*_3_. For this scenario, an index event bias is developed that explains 75% of variation in prognosis. In the fifth scenario (Sc.5), 1% of SNPs represented *G*_1_, 9% represented *G*_2_, explaining 5% and 45% of variation in incidence respectively, and 1% represented *G*_3_. The set up of this scenario developed an index event bias that explains 85% of variation in prognosis. Table I summarises the simulated scenarios. The first scenarios (Sc.1-Sc.3) show how Slope-Hunter performs, and compares its performance to the unadjusted and alternative methods [4], when the assumptions of Slope-Hunter are satisfied. Scenarios Sc.4 and Sc.5 explore performance when these underlying assumptions are not satisfied (i.e. more of the variation in incidence is explained by SNPs in class *G*_2_ than in *G*_1_).

**Table 1:** Descriptions of simulated scenarios by means of class sizes for SNPs with effects on incidence only (*G*_1_) and SNPs with effects on both incidence and prognosis (*G*_2_), and their corresponding explained variation in incidence

| Scenario | $G_1$ | | $G_2$ | | % $P$'s variance explained by confounders |
| --- | --- | --- | --- | --- | --- |
| | Size | % $I$'s variance explained | Size | % $I$'s variance explained | |
| Sc.1 | 1% | 45% | 9% | 5% | 45% |
| Sc.2 | 5% | 25% | 5% | 25% | 65% |
| Sc.3 | 1% | 25% | 9% | 25% | 65% |
| Sc.4 | 3% | 15% | 7% | 35% | 75% |
| Sc.5 | 1% | 5% | 9% | 45% | 85% |

*Abbreviations:* $I$ = incidence; $P$ = prognosis; $G_1$ = true class of SNPs with effects on incidence only; $G_2$ = true class of SNPs with effects on incidence and prognosis.
In all scenarios, 10,000 independent SNPs were simulated under Hardy-Weinberg equilibrium with minor allele frequencies drawn from a uniform distribution over the interval $[0.01, 0.49]$ . Both incidence and prognosis were simulated as quantitative traits. The heritability was 50%, and the non-genetic confounder, $U$ , explained 40% of variation in both incidence and prognosis. SNP effects, confounders and residual variation, $\varepsilon_I$ and $\varepsilon_P$ , were drawn from normal distributions. Data were simulated for 20,000 unrelated individuals. In each scenario, an index even bias is developed from genetic and non-genetic confounders that both explain certain proportion - shown in the last column - of variation in prognosis

SNP effects on incidence were simulated independently from effects on prognosis for those SNPs that affected both traits (*G*_2_). The simulations of these scenarios were repeated with correlation between SNP effects on incidence and prognosis, whereby effects were drawn from bivariate normal distribution with a particular correlation coefficient of −0.9, −0.5, 0.5 and 0.9 for the set of SNPs with effects on both traits. These led to genome-wide genetic correlations between incidence and prognosis of −0.81, −0.45, 0.45 and 0.81 for scenarios Sc.1, Sc.3 and Sc.5; −0.45, −0.25, 0.25 and 0.45 for scenario Sc.2; and −0.63, −0.35, 0.35 and 0.63 for scenario Sc.4 respectively.

Estimated SNP effects on incidence, 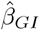 were obtained from linear regression of incidence on genotype, and the conditional estimates of SNP effects on prognosis, 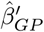 from linear regression of prognosis on genotype and incidence.

When SNP effects on incidence and prognosis are uncorrelated, the index event bias should be exactly the same for various set sizes of *G*_1_ compared with *G*_2_ as the confounding effect would be entirely due to the non-genetic component that is equally simulated across all scenarios. When there is a genetic correlation, at which the genetic component contributes to the confounding effect, the magnitude of the bias should change in a direction determined by the partial confounding effect attributed to the genetic component. For instance in our simulation set-up, when the correlation is positive, the genetic component induces bias that is in the same direction as the bias induced by the non-genetic component, both negative, resulting in a total bias of a greater magnitude than the one induced under no genetic correlation. Under negative correlations, the genetic component induces bias in the opposite direction to the non-genetic component resulting in lower total magnitudes of bias. Under the same level of genetic correlation the contribution of the genetic component to the confounding effect increases as the effect of SNPs causing both incidence and prognosis increases, e.g. from explaining 5%, as in Sc.1, to explaining 25%, as in Sc.2, of variation in incidence, see values of simulated bias at different scenarios shown in Table 2.

**Table 2:** Means and standard deviations (SD) of the differences between estimated adjustment factors using H-O and Slope-Hunter methods, and the true index event bias (*B*) over 1000 simulations of 10,000 independent SNPs, conditional on incidence as a quantitative trait, for five scenarios (Sc.1 - Sc.5) described in the first and second rows

| G. cor | Method | % $G_1$ SNPs (proportion of variation in incidence explained by $G_1$) Vs. % $G_2$ SNPs (proportion of variation in incidence explained by $G_2$) | | | | | | | | | |
| --- | --- | --- | --- | --- | --- | --- | --- | --- | --- | --- | --- |
|  |  | Sc.1: 1% (0.45) Vs. 9% (0.05) |  | Sc.2: 5% (0.25) Vs. 5% (0.25) |  | Sc.3: 1% (0.25) Vs. 9% (0.25) |  | Sc.4: 3% (0.15) Vs. 7% (0.35) |  | Sc.5: 1% (0.05) Vs. 9% (0.45) |  |
| | | Mean diff. (SD) | Mean $B$ (SD) | Mean diff. (SD) | Mean $B$ (SD) | Mean diff. (SD) | Mean $B$ (SD) | Mean diff. (SD) | Mean $B$ (SD) | Mean diff. (SD) | Mean $B$ (SD) |
| -0.90 | H-O | -0.08 (0.01) | -0.33 (0.01) | -0.43 (0.02) | -0.17 (0.01) | -0.43 (0.02) | -0.17 (0.01) | -0.62 (0.02) | -0.08 (0.02) | -0.81 (0.03) | 0.01 (0.03) |
|  | SH | -0.01 (0.02) |  | 0.01 (0.06) |  | -0.12 (0.30) |  | -0.45 (0.51) |  | -0.89 (0.07) |  |
| -0.50 | H-O | -0.04 (0.01) | -0.35 (0.01) | -0.24 (0.02) | -0.26 (0.01) | -0.23 (0.02) | -0.26 (0.01) | -0.33 (0.02) | -0.22 (0.02) | -0.43 (0.03) | -0.17 (0.03) |
|  | SH | -0.01 (0.01) |  | 0.02 (0.08) |  | 0.05 (0.12) |  | 0.02 (0.07) |  | 0.14 (0.22) |  |
| Zero | H-O | 0 (0.01) | -0.37 (0.01) | 0 (0.02) | -0.37 (0.01) | 0 (0.02) | -0.37 (0.01) | 0 (0.02) | -0.37 (0.02) | 0 (0.03) | -0.37 (0.03) |
|  | SH | 0 (0.002) |  | 0.02 (0.02) |  | 0 (0.01) |  | 0.04 (0.11) |  | 0.37 (0.12) |  |
| 0.50 | H-O | 0.05 (0.01) | -0.39 (0.01) | 0.23 (0.02) | -0.47 (0.01) | 0.23 (0.02) | -0.48 (0.01) | 0.31 (0.02) | -0.51 (0.02) | 0.40 (0.03) | -0.55 (0.03) |
|  | SH | 0 (0.003) |  | 0.02 (0.07) |  | 0 (0.04) |  | 0.13 (0.16) |  | 0.45 (0.17) |  |
| 0.90 | H-O | 0.08 (0.01) | -0.41 (0.01) | 0.40 (0.02) | -0.55 (0.01) | 0.40 (0.02) | -0.56 (0.01) | 0.55 (0.02) | -0.62 (0.02) | 0.70 (0.03) | -0.69 (0.03) |
|  | SH | 0 (0.004) |  | -0.03 (0.02) |  | 0.28 (0.40) |  | 0.72 (0.26) |  | 0.81 (0.05) |  |

*Abbreviations:* $G_1$ = true class of SNPs with effects on incidence only; $G_2$ = true class of SNPs with effects on incidence and prognosis; G. cor = genetic correlation of SNP effects on incidence and prognosis; H-O = estimator of the method of Dudbridge et al (2019) with Hedges-Olkin adjustment for regression dilution; SH = 'Slope-Hunter' estimator; Mean diff. = mean of differences between estimated adjustment factor using corresponding estimator and the true index event bias; SD = standard deviation; $B$ = true index event bias.

If a group of SNPs in the *G*_2_ class affect incidence and prognosis only through a common exposure, *E*, as depicted in Figure 3, then their genetic effects on incidence should perfectly correlate with their effects on prognosis, leading to constant proportional relationship for their SNP-prognosis to SNP-incidence associations. In this case, if the class *G*_2_ explains more of the variation in incidence than the class *G*_1_, the Slope-Hunter may be severely biased because it would completely swap the classes rather than having affordable misclassification error as expected when its assumptions are slightly violated. We simulated a sixth scenario (Sc.6) introducing this case, whereby 5% of SNPs represented the class *G*_1_, 10% represented *G*_2_, explaining 25% and 50% of variation in incidence respectively, and 5% representing *G*_3_. The class *G*_2_ was dominated by a subset of SNPs (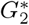: 70% of *G*_2_ that explained 35% of the total variation of incidence) whose effects on incidence and prognosis traits occur only via a common exposure. For a SNP 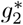 from the subset 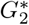, the ratio of SNP-prognosis to SNP-incidence associations can be expressed as follows:

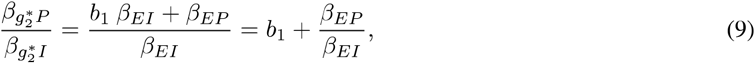

where *b*_1_ is the true confounding effect between incidence and prognosis, whereas *β*_*EI*_ and *β*_*EP*_ are the effects of exposure on incidence and prognosis respectively. This suggests a fixed ratio of SNP-prognosis to SNP-incidence associations for the group 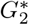, assuming small individual genetic effects as typically applies for polygenic traits. The remaining SNPs in the *G*_2_ class were simulated to have uncorrelated genetic effects on incidence and prognosis.

We performed 1000 simulations for each scenario and reported mean of differences between estimated adjustment factors and the true index event bias. The type-1 error rates, at *p* < 0.05, of SNP effects on prognosis were evaluated. Since the index event bias is proportional to the effect on incidence, as shown in equation 5, type-1 error rates vary among SNPs with different effects on incidence. Therefore, we estimated: the mean type-1 error over all SNPs with no effect on prognosis (i.e. true classes of *G*_1_ and *G*_4_); the mean type-1 error over SNPs with effects on incidence only (i.e. true class of *G* _1_) because *G* _4_ has no index event bias and its SNPs can dominate *G* _1_, when combined, due to class sizes. We estimated the family-wise type-1 error over the true class of *G*_1_, as the proportion of simulations in which at least one variant had *p* < 0.05 after Bonferroni multiple-testing adjustment for the number of SNPs. The mean power over all SNPs with effects on prognosis (true classes of *G*_2_ and *G*_3_), and over SNPs with effects on both incidence and prognosis (true classes of *G*_2_) were estimated. The mean absolute bias and mean square error across all SNPs, and across SNPs with effects on incidence (true classes of *G*_1_ and *G*_2_) were estimated.

Results of the Slope-Hunter (SH estimator) method were compared with the unadjusted estimator 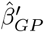 estimator of the method of Dudbridge et al (2019) with Hedges-Olkin adjustment (H-O estimator) and with simulation extrapolation adjustment (SIMEX estimator) for regression dilution [4]. Because H-O and SIMEX results were almost identical, we only presented H-O results. Furthermore, the individual SNP with highest type-1 error for the unadjusted estimator is identified and compared with the type-1 error of estimators of the adjustment methods. We identified SNPs with the greatest increase and decrease in power between the unadjusted estimator and all estimators of the compared adjustment methods, SH and H-O. The mean of maximum absolute bias is also compared between the unadjusted and adjusted estimators.

To evaluate Slope-Hunter’s estimations of the *G*_1_ class membership probabilities, we calculated the mean probability for the SNPs identified as the *G*_1_ class. Moreover, the misclassification error rate [18, 19] was obtained by comparing Slope-Hunter’s assignments of the SNPs to the *G*_1_ class with their true class status.

## 3 Results

### Simulation results

Table 2 shows means of differences between adjustment factors estimated using each of the compared methods, H-O ([4]) and SH (our approach), and the true index event bias across 1000 simulations at different scenarios Sc.1 - Sc.5 at various levels of genetic correlations. Our method, the SH, gave unbiased estimates of the adjustment factor when the proportion of variation in incidence explained by the class *G*_1_ was larger than (Sc.1) and equal (Sc.2) to those explained by the class *G*_2_. The corresponding H-O estimates were biased, with bias increasing as the genetic correlations became stronger. For Sc.1 and Sc.2, the type-1 error rates obtained from the unadjusted estimator as well as H-O and SH estimators were close to the nominal level, 0.05, when averaged over all SNPs, with slightly lower error rates for SH, see Tables 3 and 4. Since the majority of SNPs did not affect incidence, they did not suffer from index event bias. Among those with effects on incidence, for which there was bias, the type-1 error was inflated for the unadjusted analysis, ranging from 0.60 to 0.69 in Sc.1 and from 0.09 to 0.49 in Sc.2. For the H-O estimator, the type-1 error of SNPs affecting incidence was also inflated, ranging 0.08 to 0.14 in Sc.1 and from 0.15 to 0.35 for Sc.2 under genetic correlations. The type-1 error rate for our approach was consistently close to the correct rate, 0.05 - 0.06, even in the presence of genetic correlation between incidence and prognosis in both scenarios Sc.1 and Sc.2. The type-1 error for the individual SNP with the highest error under the unadjusted analysis was high, 1 in Sc.1 and 0.43 - 1 in Sc.2, but was substantially reduced using our procedure under all levels of genetic correlations, achieving the correct rate, 0.05, in Sc.1, and ranging from 0.06 to 0.14 in Sc.2. The H-O failed to reduce the error rate except under no genetic correlation. There was a similar pattern for the family-wise error rate. Overall, there were small to moderate drops in power for all adjustment methods compared to the unadjusted analysis, except under strong positive correlation where there was an increase. Our method consistently achieved higher power rates than the H-O method under all levels of genetic correlations. For some individual SNPs the power rate was low under the unadjusted analysis but was substantially increased under all adjustment approaches, with the greatest increase under the SH method when there was positive correlation between incidence and prognosis. Under negative correlation, the H-O method had greater increase in power for some individual SNPs compared to SH method, and similar increase when there was no genetic correlation. Our procedure had the lowest absolute bias compared to the unadjusted as well as the other adjusted analysis under all levels of genetic correlations. There were similar patterns for the mean square error.

**Table 3:**
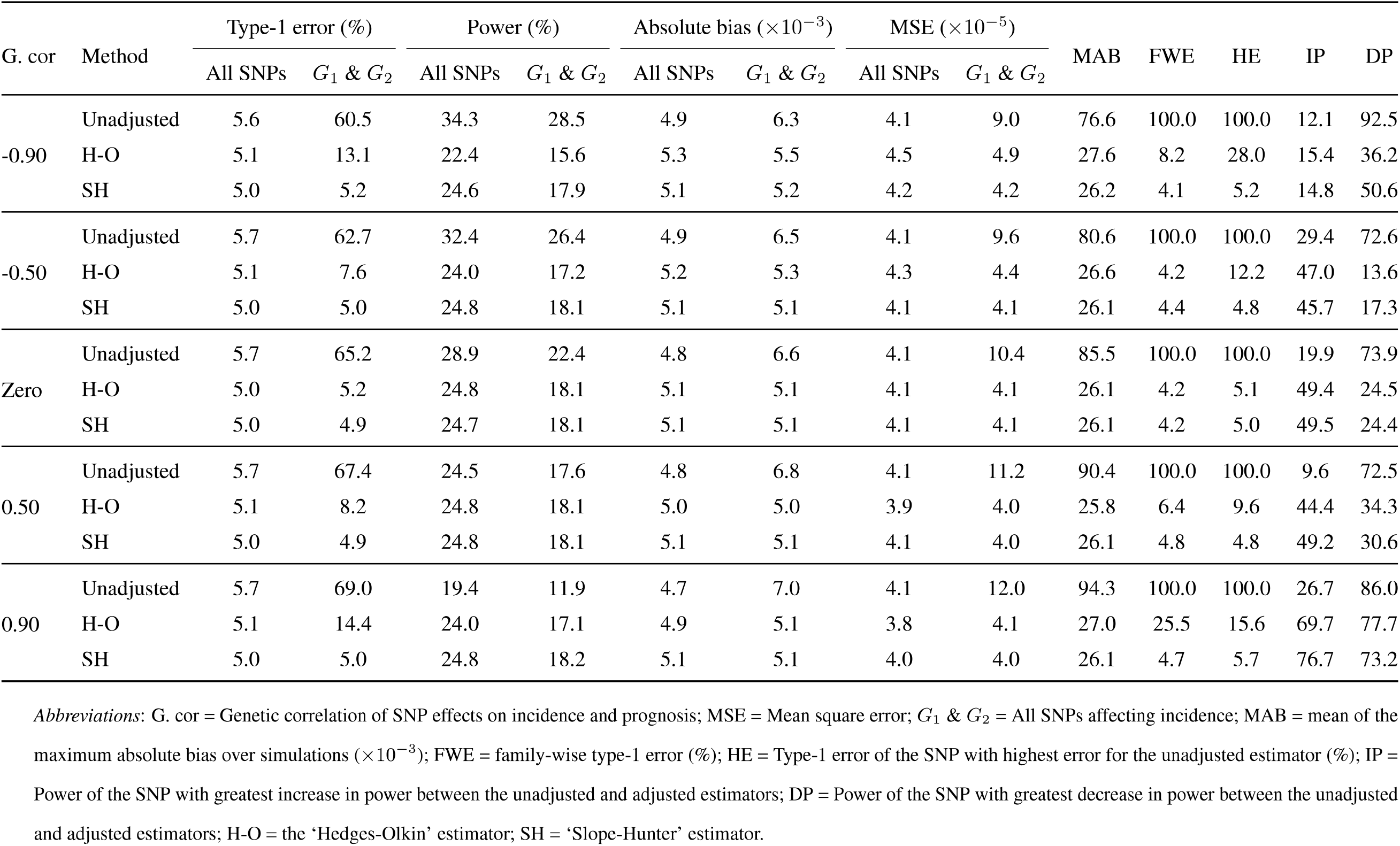
Type-1 error and power at *p* < 0.05, absolute bias and mean square error over 1000 simulations of 10,000 independent SNPs, conditional on incidence as a quantitiative trait, for Sc.1: 1% of SNPs have effects on incidence only (explaining 45 % of its variation), 1% on prognosis only and 9% on both incidence and prognosis (explaining 5% of variation in incidence). Heritability of incidence and prognosis is 50% with the genetic correlation between SNP effects on incidence and prognosis shown in the first column. Non-genetic common factors explain 40% of variation in both incidence and prognosis. The index event bias explains ~ 45 % of variation in prognosis

**Table 4:** Type-1 error and power at *p* < 0.05, absolute bias and mean square error over 1000 simulations of 10,000 independent SNPs, conditional on incidence as a quantitative trait, for Sc.2: 5% of SNPs have effects on incidence only (explaining 25% of its variation), 5% on prognosis only and 5% on both incidence and prognosis (explaining 25% of variation in incidence). Heritability of incidence and prognosis is 50% with the genetic correlation between SNP effects on incidence and prognosis shown in the first column. Non-genetic common factors explain 40% of variation in both incidence and prognosis. The index event bias explains~ 65% of variation in prognosis

| G. cor | Method | Type-1 error (%) | | | Power (%) | | | Absolute bias ($\times 10^{-3}$) | | | MSE ($\times 10^{-5}$) | | | MAB | FWE | HE | IP | DP |
| --- | --- | --- | --- | --- | --- | --- | --- | --- | --- | --- | --- | --- | --- | --- | --- | --- | --- | --- |
| | | All SNPs | $G_1$ & $G_2$ | | All SNPs | $G_1$ & $G_2$ | | All SNPs | $G_1$ & $G_2$ | | All SNPs | $G_1$ & $G_2$ | | | | | | |
| -0.90 | Unadjusted | 5.2 | 8.9 |  | 61.0 | 62.6 |  | 5.3 | 5.9 |  | 4.4 | 5.4 |  | 27.0 | 8.8 | 43.4 | 12.7 | 100.0 |
|  | H-O | 5.9 | 21.8 |  | 42.8 | 32.9 |  | 6.6 | 10.2 |  | 7.1 | 16.2 |  | 43.6 | 89.8 | 97.0 | 63.5 | 5.7 |
|  | SH | 5.0 | 5.3 |  | 58.2 | 57.5 |  | 5.3 | 5.4 |  | 4.4 | 4.6 |  | 26.9 | 5.9 | 7.3 | 26.4 | 97.4 |
| -0.50 | Unadjusted | 5.6 | 15.1 |  | 62.2 | 64.1 |  | 5.2 | 6.7 |  | 4.2 | 7.0 |  | 29.3 | 37.7 | 80.8 | 6.9 | 99.6 |
|  | H-O | 5.4 | 11.5 |  | 52.6 | 49.8 |  | 5.9 | 7.3 |  | 5.5 | 8.3 |  | 31.9 | 13.6 | 60.1 | 100.0 | 8.8 |
|  | SH | 5.1 | 5.9 |  | 58.6 | 58.3 |  | 5.2 | 5.4 |  | 4.3 | 4.6 |  | 26.5 | 5.6 | 14.5 | 83.3 | 73.9 |
| Zero | Unadjusted | 6.2 | 25.8 |  | 61.5 | 60.8 |  | 5.0 | 8.1 |  | 4.1 | 10.3 |  | 37.6 | 99.5 | 99.4 | 5.1 | 95.8 |
|  | H-O | 5.0 | 5.0 |  | 57.6 | 56.3 |  | 5.1 | 5.2 |  | 4.1 | 4.2 |  | 25.7 | 5.0 | 4.6 | 97.3 | 7.9 |
|  | SH | 5.0 | 5.0 |  | 57.8 | 56.5 |  | 5.1 | 5.1 |  | 4.0 | 4.2 |  | 25.5 | 5.1 | 5.7 | 95.8 | 7.2 |
| 0.50 | Unadjusted | 6.9 | 38.2 |  | 60.7 | 56.5 |  | 4.8 | 9.9 |  | 4.0 | 15.3 |  | 41.6 | 100.0 | 100.0 | 4.4 | 98.3 |
|  | H-O | 5.5 | 14.7 |  | 59.1 | 55.2 |  | 4.6 | 6.4 |  | 3.4 | 6.4 |  | 27.8 | 34.1 | 77.3 | 75.3 | 43.9 |
|  | SH | 5.1 | 6.0 |  | 57.0 | 54.7 |  | 5.0 | 5.2 |  | 3.9 | 4.4 |  | 25.4 | 7.5 | 9.9 | 97.4 | 8.5 |
| 0.90 | Unadjusted | 7.4 | 48.6 |  | 53.8 | 40.0 |  | 4.5 | 11.4 |  | 4.1 | 20.7 |  | 49.3 | 100.0 | 100.0 | 5.9 | 77.9 |
|  | H-O | 6.7 | 34.9 |  | 56.6 | 46.3 |  | 4.4 | 8.9 |  | 3.4 | 12.6 |  | 39.4 | 100.0 | 100.0 | 31.5 | 56.0 |
|  | SH | 5.0 | 5.1 |  | 57.0 | 55.8 |  | 5.0 | 5.1 |  | 3.9 | 4.1 |  | 25.3 | 5.0 | 5.6 | 98.9 | 6.1 |

*Abbreviations:* G. cor = Genetic correlation of SNP effects on incidence and prognosis; MSE = Mean square error; $G_1$ & $G_2$ = All SNPs affecting incidence; MAB = mean of the maximum absolute bias over simulations ( $\times 10^{-3}$ ); FWE = family-wise type-1 error (%); HE = Type-1 error of the SNP with highest error for the unadjusted estimator (%); IP = Power of the SNP with greatest increase in power between the unadjusted and adjusted estimators; DP = Power of the SNP with greatest decrease in power between the unadjusted and adjusted estimators; H-O = the 'Hedges-Olkin' estimator; SH = 'Slope-Hunter' estimator.

All methods consistently performed less well as the proportion of variation in incidence explained by the *G*_1_ class decreased in the presence of genetic correlation. However, the SH estimator consistently outperformed the other estimators providing the closest estimates to the true bias and the lowest type-1 error rates, see Table 2 and Tables 5-7. Under no genetic correlation, the SH method performed as well as the H-O method when *G*_1_ explains at least as much variation in incidence as *G*_2_ as in Sc.1 - Sc.3, see Tables 2 - 5. In Sc.3, the SH method consistently outperformed the other methods - in terms of bias and type-1 error - under all levels of genetic correlation. SH performed less well under strong genetic correlation in Sc.3 compared to Sc.1 (i.e. performed less well where the variance of incidence explained by the *G*_1_ class was lower). SH performed less well under strong genetic correlation in Sc.3 compared to Sc.2 (i.e. performed less well where the number of SNPs in the *G*_1_ class was lower). Tables 6 and 7 show results of type-1 error, power, absolute bias and mean square error for Sc.4 and Sc.5 respectively. As expected, the SH estimator was more biased as the class *G*_1_ explained less variation in *I* than the class *G*_2_, with worse performance under stronger genetic correlation. But, the bias in the SH estimator was lower than in H-O estimator for small and moderate index event bias (ranging from - 0.5 to 0). Our approach provided lower type-1 error rates than the unadjusted analyses as well as the H-O method except under strong correlation in Sc.4. The SH estimator provided equivalent power, compared with the alternatives, except under strong correlation, where it had slightly lower power in Sc. 4 and Sc.5. The absolute bias and mean square errors showed similar pattern to the results of type-1 error. In Sc.6, the SH estimator was severely biased providing worse type-1 error rates, 0.56 versus 0.36, with lower power, 0.25 versus 0.45, compared with the unadjusted estimator, see Table S1 in the supplementary material.

**Table 5:**
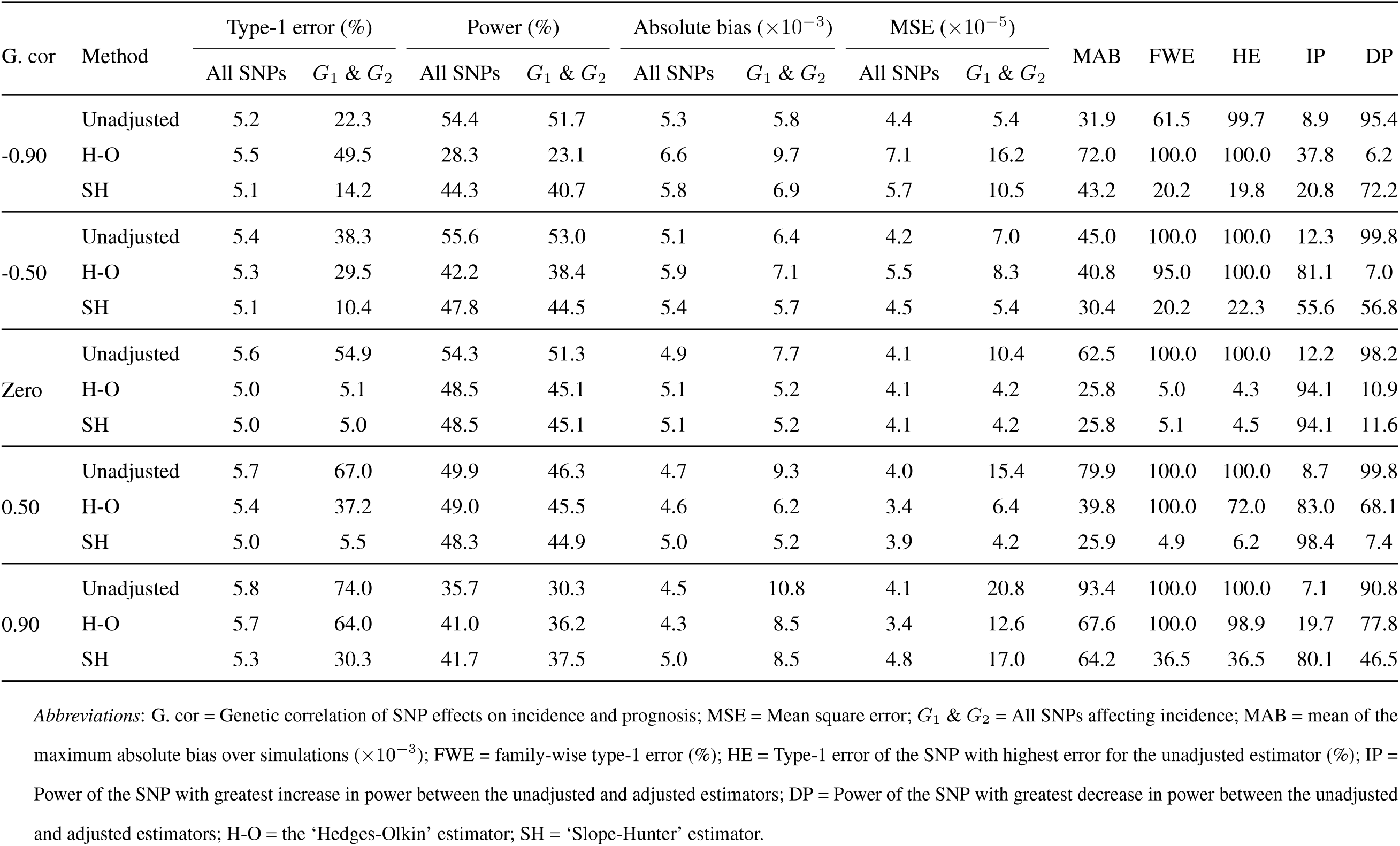
Type-1 error and power at *p* < 0.05, absolute bias and mean square error over 1000 simulations of 10,000 independent SNPs, conditional on incidence as a quantitiative trait, for Sc.3: 1% of SNPs have effects on incidence only (explaining 25% of its variation), 1% on prognosis only and 9% on both incidence and prognosis (explaining 25% of variation in incidence). Heritability of incidence and prognosis is 50% with the genetic correlation between SNP effects on incidence and prognosis shown in the first column. Non-genetic common factors explain 40% of variation in both incidence and prognosis. The index event bias explains ~ 65% of variation in prognosis

**Table 6:**
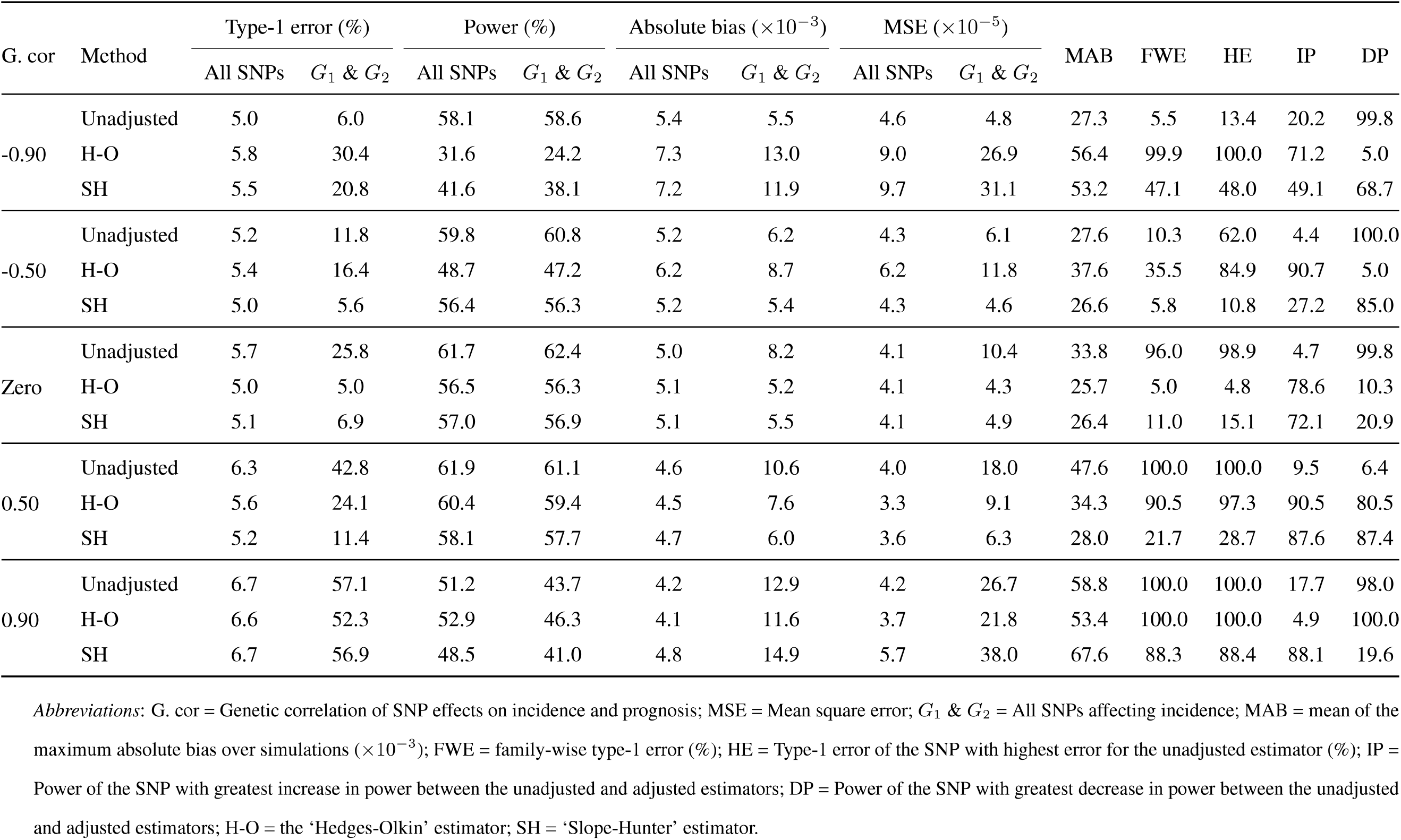
Type-1 error and power at *p* < 0.05, absolute bias and mean square error over 1000 simulations of 10,000 independent SNPs, conditional on incidence as a quantitiative trait, for Sc.4: 3% of SNPs have effects on incidence only (explaining 15% of its variation), 3% on prognosis only and 7% on both incidence and prognosis (explaining 35% of variation in incidence). Heritability of incidence and prognosis is 50% with the genetic correlation between SNP effects on incidence and prognosis shown in the first column. Non-genetic common factors explain 40% of variation in both incidence and prognosis. The index event bias explains ~ 75% of variation in prognosis

**Table 7:**
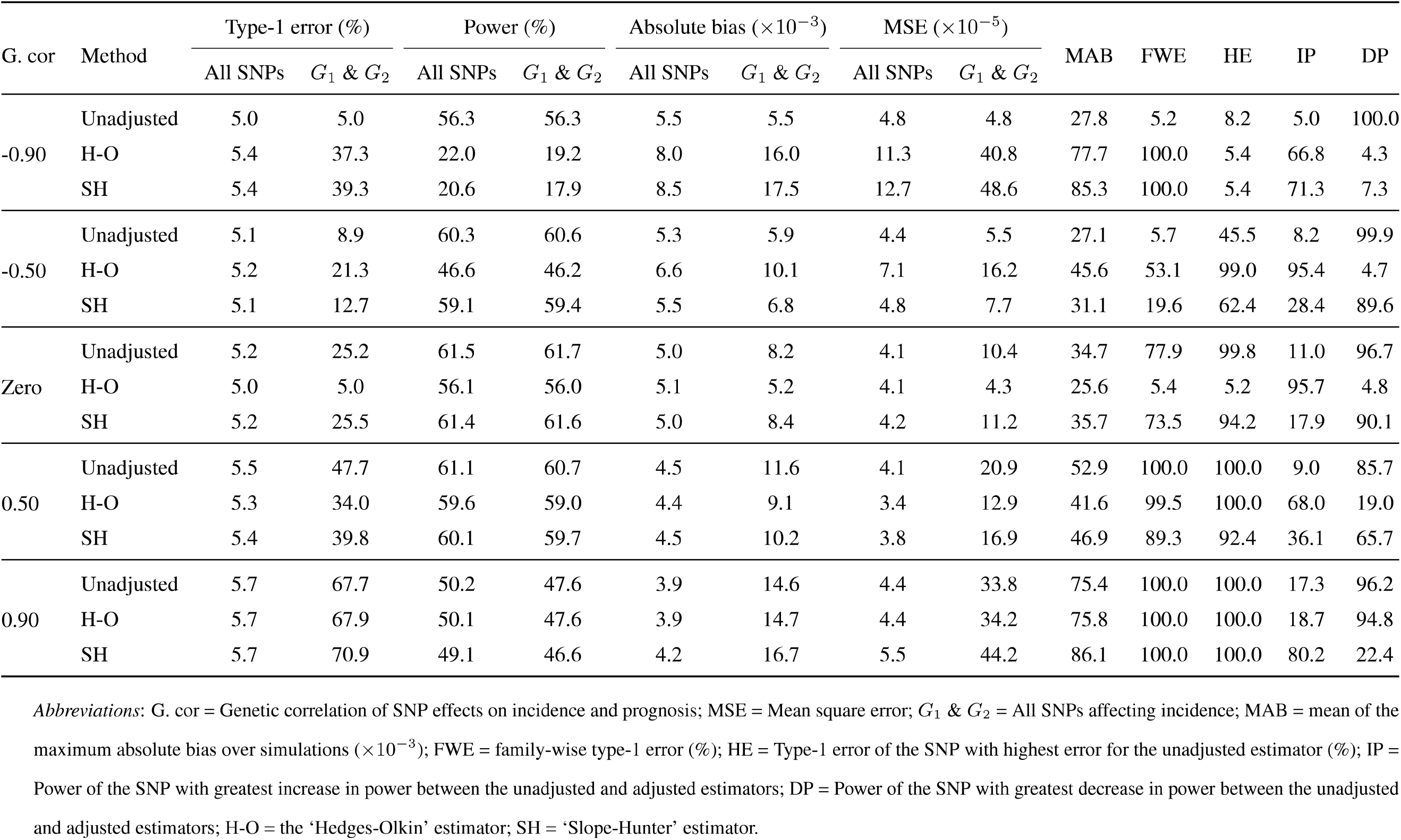
Type-1 error and power at *p* < 0.05, absolute bias and mean square error over 1000 simulations of 10,000 independent SNPs, conditional on incidence as a quantitiative trait, for Sc.5: 1% of SNPs have effects on incidence only (explaining 5% of its variation), 1% on prognosis only and 9% on both incidence and prognosis (explaining 45% of variation in incidence). Heritability of incidence and prognosis is 50% with the genetic correlation between SNP effects on incidence and prognosis shown in the first column. Non-genetic common factors explain 40% of variation in both incidence and prognosis. The index event bias explains ~ 85% of variation in prognosis

The mean probabilities of memberships assignments to the cluster *G*_1_ obtained by the SH were similar, ranging between 0.80 to 0.90, in Sc.1 and Sc.3. The classification error rates were slightly lower in Sc.3 compared with Sc.1, reflecting better identification of variants belonging to the true *G*_1_ class, see Table S2 in the supplementary material. Figures 8 and 9 present uncertainty plots for the variants assigned to the *G*_1_ cluster by the SH method in four simulations selected randomly from the 1000 simulations of Sc.l with - 0.9 and 0.9 correlations respectively. The misclassified SNPs, indicated by vertical black lines, were the ones with the highest uncertain identification. Figure S1 in the supplementary material shows uncertainty plots for the Sc.6, in which almost all variants of the *G*_1_ cluster were misclassified regardless of their uncertainty levels.

**Figure 8:**
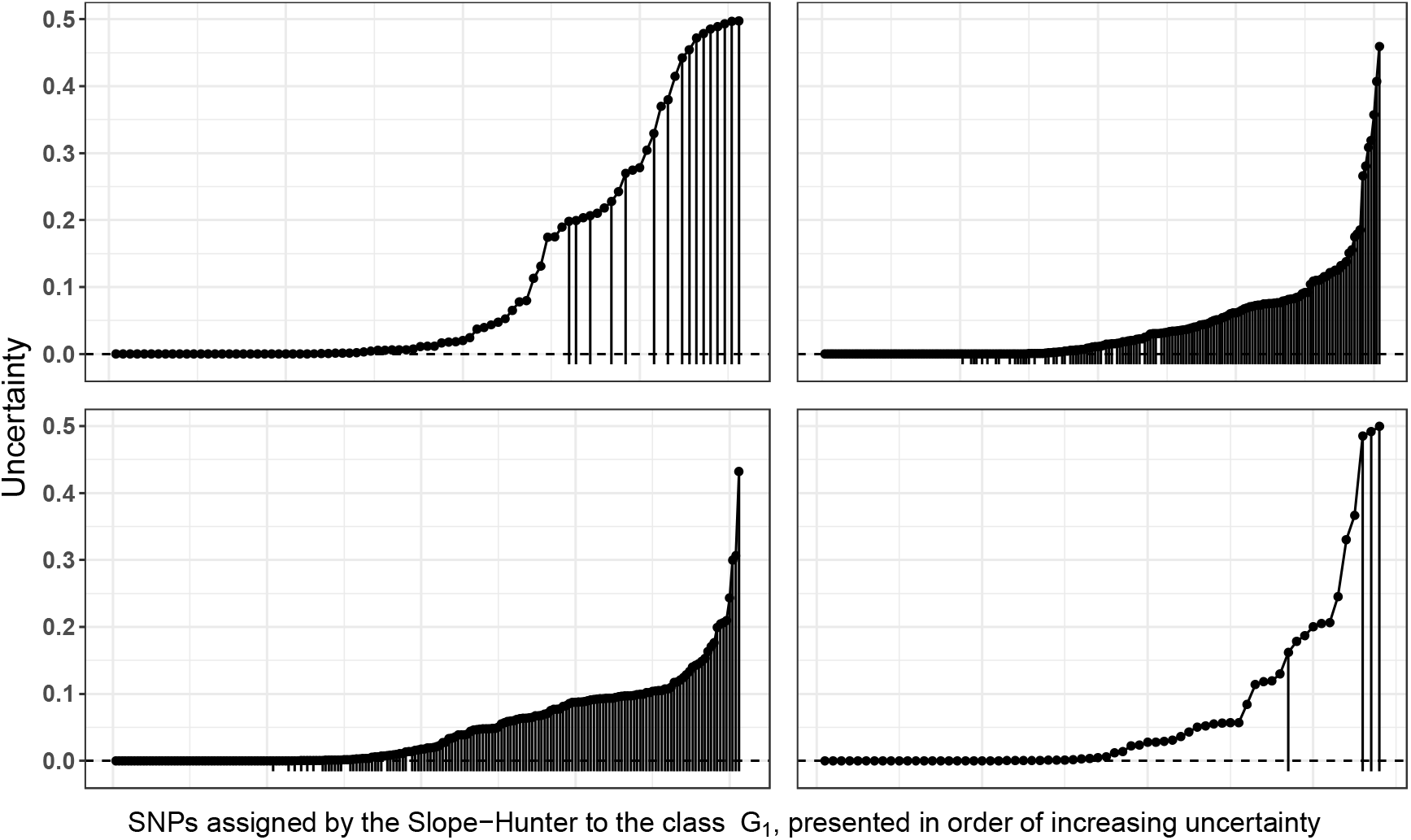
Uncertainty plot for SNPs identified by the Slope-Hunter method as the variants affecting incidence only from four simulations of scenario 4 in which 1% of SNPs have effects on incidence only and 9% on both incidence and prognosis with a correlation of −0.9, explaining 45% and 5% respectively of variation in incidence. The vertical lines indicate misclassified SNPs.

**Figure 9:**
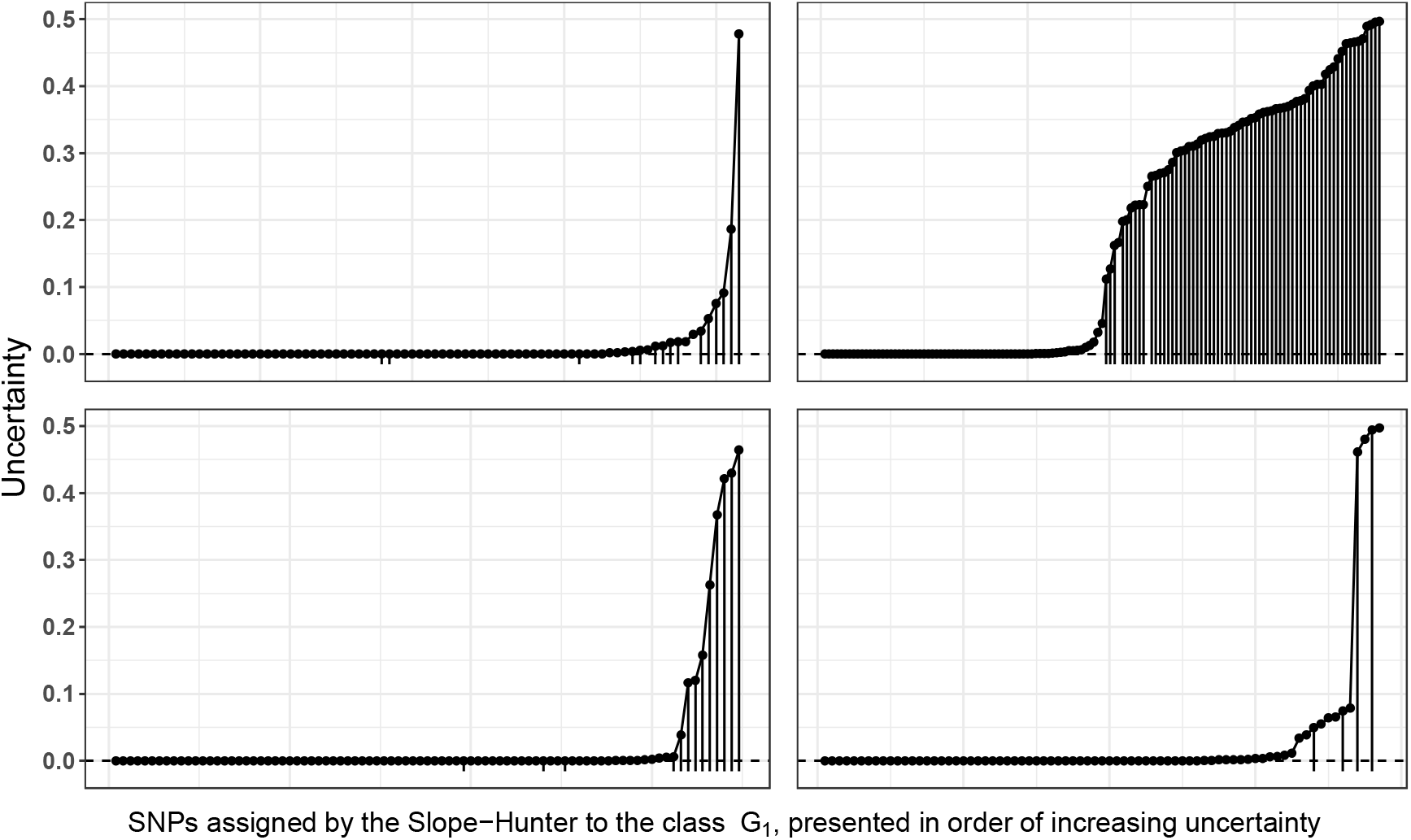
Uncertainty plot for SNPs identified by the Slope-Hunter method as the variants affecting incidence only from four simulations of scenario 4 in which 1% of SNPs have effects on incidence only and 9% on both incidence and prognosis with a correlation of 0.9, explaining 45% and 5% respectively of variation in incidence. The vertical lines indicate misclassified SNPs.

## 4 Discussion

Analysis of causal effects on prognosis, such as subsequent disease events, severity and survival time, is increasingly motivated by many large collections of GWAS for disease cases. Such case-only studies are liable to index event bias, whereby independent causes of the incidence become correlated when selecting only on cases and then may confound analysis of prognosis. We have proposed an approach that overcomes a major disadvantage of previous methods, and showed that it provides unbiased estimates of SNP-prognosis associations in a variety of situations including the presence of genetic correlations between incidence and prognosis. Our approach aims to identify the set of SNPs with effects only on incidence and uses it to estimate and adjust for the index event bias induced by the confounder effects. Therefore, our approach is robust against the violation of the ‘InCLUDE’ assumption, that is the direct genetic effects on prognosis are assumed to be linearly uncorrelated with effects on incidence. Our analytic approach assumes the analysed SNPs are independent, do not interact with the confounders, and have linear effects with the incidence and prognosis. Moreover, it assumes that the variance of incidence explained by the class *G*_1_ is at least as much as that explained by the class *G*_2_. Under satisfaction of these underlying assumptions, our procedure is highly adaptive in dealing with genetic correlations, and can maintain excellent trade-off between type-1 error rates and power and produce lower mean square error compared to the other methods. Independence of the analysed SNPs can be achieved by selecting from GWAS through LO-pruning prioritising by p-values of SNP-incidence associations. However, if traits are generated under non-linear models or the variance of incidence explained by the SNPs affecting only incidence is extremely small, a biased adjustment factor might be derived by our procedure leading to inexact correction for the index event bias.

We simulated an index event bias by analysing prognosis conditional on a continuous incidence trait. Dudbridge et. al. (2018) showed that similar index event bias is induced when incidence trait is a binary disease and prognosis is analysed in cases only [4]. Our simulations showed that our approach consistently achieved the minimum type-1 error rates, with slightly less power on average and considerably higher power for some individual SNPs, compared with the unadjusted analysis. Compared with alternative methods, our procedure had lower type-1 error rates and higher average power under various levels of genetic correlations between incidence and prognosis. All methods had worse type-1 error rates under genetic correlations as the proportion of incidence variance explained by SNPs affecting incidence only reduced. However our approach had better type-1 error rates than the alternatives when this proportion was not very small, except under strong correlations where it produced worse type-1 error than the unadjusted analysis but better type-1 error and less biased estimates than the compared methods.

Our method relies on obtaining an estimate of the bias adjustment factor using information from the SNPs that only affect incidence, the *G* _1_ class. Since identification of this class is more accurate when its variants are more concentrated around the cluster’s slope [12], then our procedure produced biased estimates as the SNPs in this group explained less of variation in incidence than the variants affecting both incidence and prognosis. The unadjusted analysis similarly performed worse as the proportion of incidence’s variance explained by SNPs affecting only incidence was decreased, while the corresponding proportion for the SNPs affecting both incidence and prognosis was increased. This could be due to the increase in confounder effects, and hence the bias, resulting from increased effects of SNPs influencing both incidence and prognosis, the *G*_2_ class. Increasing effects of SNPs in the *G*_2_ class on incidence in the presence of genetic correlation between incidence and prognosis caused more violation of the ‘InCLUDE’ assumption, a critical assumption for the H-O estimator [4], leading to more biased estimates of these methods. The SH estimator may provide less good performance in index event bias correction when the misclassification in the *G*_1_ cluster severely influences its pattern. This may occur if the variants of *G*_2_, that are misclassified to the *G*_1_ cluster, explain more variation in incidence than the class *G*_1_ (as in Sc.4 and Sc.5); or explain equal variation in *I* but the size of the *G*_1_ cluster is small, as in Sc.3, such that a misclassified SNP would have a bigger effect on its pattern. In either case, the influence of misclassification is more pronounced under stronger genetic correlation, which applies only, by definition, to SNPs affecting both incidence and prognosis.

Our procedure requires user choices for the input parameters, Λ, *η* and *δ*. These choices impact identification of the class of SNPs with effects only on incidence, *G*_1_. For example, retaining a sub-sample of only l % of SNPs with no effects on neither incidence nor prognosis, the *G*_4_ class, may not be sufficient to aid pattern recognition of the *G*_1_ cluster in the dimensional space. Therefore, it may be beneficial to retain larger sub-sample of the class *G*_4_, but too large sub-samples may distort pattern of the class *G*_1_ leading to poor cluster identification. Fitting cluster-based models iteratively using a number of proportions given by the set A allows users to specify many potential sub-sample sizes to be retained in the analysis and the algorithm can then pick the best size as the one leading to the highest precision of estimate of the adjustment factor. Inclusion of more SNPs is the final stage of analysis as potential members of either the class *G*_1_ or *G*_2_, when more lenient *p*-value threshold, *η*, is used, could lead to improving efficiency due to potential increase in underlying SNPs of *G*_1_ class. But, it might also result in including a fraction of SNPs with no effects on incidence in the *G*_1_ class, whose pattern may be obscured if this fraction is large. Under the assumptions of our procedure, the SNPs in *G*_2_ class are expected to be scattered around the slope, *b*_1_, with higher variations than the SNPs in *G*_1_ class. Then, a valid cluster solution should have a relatively small Euclidean distance between the means of clusters *G*_1_ and *G*_2_. Our simulations showed that this distance could be maximally equal to the minimum of standard deviations of both clusters on each dimension. However, the parameter *δ* is used as a multiplier of the minimum standard deviation that can be set by the user to define valid cluster solutions for different data structures.

The main idea of our procedure can be exploited in future in the context of the MR analysis using a large number of genetic variants including invalid instruments, particularly for experiments in which effects of instruments on exposure and outcome are correlated[20]. This potential direction may be beneficial in robustly estimating causal effects, checking violation of MR assumptions, and providing probabilistic identification of the valid instruments in a given problem. A few methods have been recently developed with a conceptual similarity to the Slope-Hunter in the context of MR analysis, that is they aim to identify the valid instruments, and then use this class of genetic variants to estimate causal effect of an exposure on outcome. These include the MR-mix [17] and CAUSE [21] methods. The Slope-Hunter approach can be adapted in future to be used in conjunction with these methods to form a consensus results of GWAS hits for a trait since MR assumptions can not be exactly verified. For instance, the MR-mix method relies on the ZEMPA assumption implying that the number of valid instruments are assumed to be larger than any other group sizes of invalid instruments with unique estimates of the causal effect. The Slope-Hunter has a less restrictive assumption implying that the group of SNPs affecting incidence only explains at least as much variation in incidence as the group of SNPs affecting both incidence and prognosis.

Our study has several strengths. It provides a novel framework that uses a cluster-based model approach to correct for index event bias even in the presence of genetic correlations between incidence and prognosis. Our approach provides estimates for the membership probability of each SNP to each cluster allowing us to evaluate the obtained cluster solutions. We evaluated our developed approach in a variety of situations including different levels of genetic correlations, leading to different magnitudes of genetic confounders, and different number of SNPs with effects only on incidence. Our study compared the performance of the SH method with the unadjusted analysis as well as with other alternative methods in terms of many statistical criteria including type-1 error, power, bias and mean squared error. On the other hand, there are a number of limitations. Although we have examined the SH performance in a wide range of situations, including violating the ‘InCLUDE’ assumption and larger effects for the *G*_2_ class, we have not examined the sensitivity to non-linearity or to interaction between confounder and variant’s effects. We have not examined sensitivity to the user-set parameters, and these may be refined in light of how the method performs in practice. There is not a single criterion for validity of the slope-hunter approach, as it will depend on how separated the classes *G*_1_ and *G*_2_ are and whether the class *G*_1_ is correctly identified. However, the performance of the SH is influenced by the identified pattern of the *G*_1_ class, and hence its slope, rather than the exact accuracy of classifying the *G*_1_ variants. Therefore, the SH can afford a reasonable amount of misclassification error as long as the misclassified SNPs do not severely distort the estimated pattern of the target class. Assumptions of the Slope-Hunter are not testable, but the estimated probabilities of fitted cluster solutions for *G*_1_ SNPs can be used for diagnosis purposes to assess the method performance.

We have proposed an approach for adjusting for index event bias in GWAS of subsequent events even in the presence of genetic correlation between incidence and prognosis. We recommend that this approach is used in GWAS of events after incidence of a disease, to minimise the bias due to conditioning on incidence. This approach is also recommended for subsequent use of GWAS results, such as in MR analyses of the effect of exposure on prognosis. All procedures described in this manuscript have been implemented into an open source R package named ‘Slope-Hunter’.

## References

[1] Lavinia Paternoster, Kate Tilling, and George Davey Smith. Genetic epidemiology and mendelian randomization for informing disease therapeutics: conceptual and methodological challenges. PLoS genetics, 13(10):e1006944, 2017.

[2] James Yarmolinsky, Kaitlin H Wade, Rebecca C Richmond, Ryan J Langdon, Caroline J Bull, Kate M Tilling, Caroline L Relton, Sarah J Lewis, George Davey Smith, and Richard M Martin. Causal inference in cancer epidemiology: what is the role of mendelian randomization? Cancer Epidemiology and Prevention Biomarkers, 27(9):995–1010, 2018.

[3] Hanieh Yaghootkar, Michael P Bancks, Sam E Jones, Aaron McDaid, Robin Beaumont, Louise Donnelly, Andrew R Wood, Archie Campbell, Jessica Tyrrell, Lynne J Hocking, et al. Quantifying the extent to which index event biases influence large genetic association studies. Human molecular genetics, 26(5):1018–1030, 2016.

[4] Frank Dudbridge, Richard J Allen, Nuala A Sheehan, A Floriaan Schmidt, James C Lee, R Gisli Jenkins, Louise V Wain, Aroon D Hingorani, and Riyaz S Patel. Adjustment for index event bias in genome-wide association studies of subsequent events. Nature communications, 10(1):1561, 2019.

[5] Stephen R Cole and Miguel A Heman. Constructing inverse probability weights for marginal structural models. American journal of epidemiology, 168(6):656–664, 2008.

[6] Geraldine M Clarke, Kirk Rockett, Katja Kivinen, Christina Hubbart, Anna E Jeffreys, Kate Rowlands, Muminatou Jallow, David J Conway, Kalifa A Bojang, Margaret Pinder, et al. Characterisation of the opposing effects of g6pd deficiency on cerebral malaria and severe malarial anaemia. Elife, 6:e15085, 2017.

[7] Malaria Genomic Epidemiology Network, Kirk A Rockett, Geraldine M Clarke, Kathryn Fitzpatrick, Christina Hubbart, Anna E Jeffreys, Kate Rowlands, Rachel Craik, Muminatou Jallow, David J Conway, et al. Reappraisal of known malaria resistance loci in a large multicenter study. Nature genetics, 46(11):1197, 2014.

[8] James A Watson, Stije J Leopold, Julie A Simpson, Nicholas PJ Day, Arjen M Dondorp, and Nicholas J White. Collider bias and the apparent protective effect of glucose-6-phosphate dehydrogenase deficiency on cerebral malaria. eLife, 8:e43154, 2019.

[9] Chi-Fa Hung, Margarita Rivera, Nick Craddock, Michael J Owen, Michael Gill, Ania Korszun, Wolfgang Maier, Ole Mors, Martin Preisig, John P Rice, et al. Relationship between obesity and the risk of clinically significant depression: Mendelian randomisation study. The British Journal of Psychiatry, 205(1):24–28, 2014.

[10] Andrea Ganna, Samira Salihovic, Johan Sundstrom, Corey D Broeckling, Asa K Hedman, Patrik KE Magnusson, Nancy L Pedersen, Anders Larsson, Agneta Siegbahn, Mihkel Zilmer, et al. Large-scale metabolomic profiling identifies novel biomarkers for incident coronary heart disease. PLoS genetics, 10(12):e1004801, 2014.

[11] Robyn E Wootton, Rebecca C Richmond, Bobby G Stuijfzand, Rebecca B Lawn, Hannah M Sallis, Gemma MJ Taylor, Hannah J Jones, Stanley Zammit, George Davey Smith, and Marcus R Munafo. Causal effects of lifetime smoking on risk for depression and schizophrenia: Evidence from a mendelian randomisation study. Biorxiv, page 381301, 2018.

[12] Chris Fraley and Adrian E Raftery. Model-based clustering, discriminant analysis, and density estimation. Journal of the American statistical Association, 97(458):611–631, 2002.

[13] Chris Fraley and Adrian E Raftery. Bayesian regularization for normal mixture estimation and model-based clustering. Journal of classification, 24(2):155–181, 2007.

[14] Luca Scrucca, Michael Fop, T Brendan Murphy, and Adrian E Raftery. mclust 5: clustering, classification and density estimation using gaussian finite mixture models. The Rjournal, 8(1):289, 2016.

[15] Jeffrey D Banfield and Adrian E Raftery. Model-based gaussian and non-gaussian clustering. Biometrics, pages 803–821, 1993.

[16] Gilles Celeux and Gerard Govaert. Gaussian parsimonious clustering models. 1993.

[17] Guanghao Qi and Nilanjan Chatterjee. Mendelian randomization analysis using mixture models for robust and efficient estimation of causal effects. Nature communications, 10(1):1941, 2019.

[18] Osama Mahmoud, Andrew Harrison, Aris Perperoglou, Asma Gul, Zardad Khan, Metodi V Metodiev, and Berthold Lausen. A feature selection method for classification within functional genomics experiments based on the proportional overlapping score. BMC bioinformatics, 15(1):274, 2014.

[19] Osama Mahmoud, Andrew Harrison, Asma Gul, Zardad Khan, Metodi V Metodiev, and Berthold Lausen. Minimizing redundancy among genes selected based on the overlapping analysis. In Analysis of Large and Complex Data, pages 275–285. Springer, 2016.

[20] Jack Bowden, George Davey Smith, and Stephen Burgess. Mendelian randomization with invalid instruments: effect estimation and bias detection through egger regression. International journal of epidemiology, 44(2):512–525, 2015.

[21] Jean Morrison, Nicholas Knoblauch, Joe Marcus, Matthew Stephens, and Xin He. Mendelian randomization accounting for horizontal and correlated pleiotropic effects using genome-wide summary statistics. bioRxiv, page 682237, 2019.

