## Supplementary material for "Slope-Hunter: A robust method for index-event bias correction in genome-wide association studies of subsequent traits"

Table S1: Type-1 error and power at $p < 0.05$, absolute bias and mean square error over 1000 simulations of 10,000 independent SNPs, conditional on incidence as a quantitiative trait, for Sc.6: $5\%$ of SNPs have effects on incidence only ($G_1$) and $10\%$ on both incidence and prognosis ($G_2$), explaining $25\%$ and $50\%$ respectively of variation in incidence. Of the $G_2$ group, 70% affect incidence and prognosis through the same exposure, explaining 35% of variation in incidence. The rest of $G_2$ SNPs had zero genetic correlation between effects on incidence and prognosis. The heritability of incidence and prognosis is $50\%$. Non-genetic common factors explain $40\%$ of variation in both incidence and prognosis. The Mean of estimated class membership probabilities obtained by the Slope-Hunter for the $G_1$ cluster and their means of misclassification error rates are shown

| Method | Type-1 error (%) | | Power (%) | | Absolute bias ($\times 10^{-3}$) | | MSE ($\times 10^{-5}$) | |
|---|---|---|---|---|---|---|---|---|
| | All SNPs | $G_1$ & $G_2$ | All SNPs | $G_1$ & $G_2$ | All SNPs | $G_1$ & $G_2$ | All SNPs | $G_1$ & $G_2$ |
| Unadj. | 6.8 | 35.7 | 51.3 | 44.8 | 5.1 | 9.8 | 4.8 | 15.4 |
| SH | 8.0 | 55.7 | 36.1 | 24.9 | 7.0 | 17.7 | 11.0 | 50.6 |
| | MAB | FWE | HE | IP | DP | Mean diff. (SD); $B$ | Prob. | E. rate (%) |
| Unadj. | 46.3 | 100.0 | 100.0 | 7.4 | 100.0 | — | — | — |
| SH | 81.1 | 100.0 | 100.0 | 89.6 | 9.1 | 0.91 (0.11); -0.45 | 0.752 | 97.7 |

*Abbreviations*: MSE = mean square error; $G_1$ & $G_2$ = all SNPs affecting incidence; MAB = mean of the maximum absolute bias over simulations ($\times 10^{-3}$); FWE = family-wise type-1 error (%); HE = type-1 error of the SNP with highest error for the unadjusted estimator (%); Unadj. = the unadjusted analysis; SH = the Slope-Hunter estimator; IP = power of the SNP with greatest increase in power between the unadjusted and SH estimators; DP = power of the SNP with greatest decrease in power between the unadjusted and SH; Mean diff. = mean of differences between adjustment factor estimated using SH method and the true index event bias; SD = standard deviations; $B$ = true index event bias; Prob. = mean of estimated class membership probabilities obtained by the SH for the $G_1$ cluster; E. rate = mean of misclassification error rates for the SNPs assigned to the $G_1$ cluster by the SH method

Table S2: Means of estimated class membership probabilities obtained by the Slope-Hunter for the $G_1$ cluster and their means of misclassification error rates over 1000 simulations of 10,000 independent SNPs, conditional on incidence as a quantitative trait, for Sc.1 and Sc.3, with higher and lower proportions of variation in incidence explained by the $G_1$ class respectively, under different levels of genetic correlation of SNP effects on incidence and prognosis

| | % $G_1$ SNPs ($V(I)$ explained by $G_1$) Vs. % $G_2$ SNPs ($V(I)$ explained by $G_2$) | | | |
|---|---|---|---|---|
| | **Sc.1**: 1% (0.45) Vs. 9% (0.05) | | **Sc.3**: 1% (0.25) Vs. 9% (0.25) | |
| G. cor | Mean probability | Mean error rate (%) | Mean probability | Mean error rate (%) |
| -0.90 | 0.860 | 37.0 | 0.870 | 23.0 |
| -0.50 | 0.855 | 47.0 | 0.850 | 27.0 |
| Zero | 0.859 | 50.0 | 0.820 | 21.0 |
| 0.50 | 0.852 | 46.0 | 0.800 | 23.0 |
| 0.90 | 0.847 | 40.0 | 0.900 | 39.0 |

*Abbreviations*: $V(I)$ = explained variation of incidence (as a proportion); $G_1$ = true class of SNPs with effects on incidence only; $G_2$ = true class of SNPs with effects on incidence and prognosis; G. cor = genetic correlation of SNP effects on incidence and prognosis.

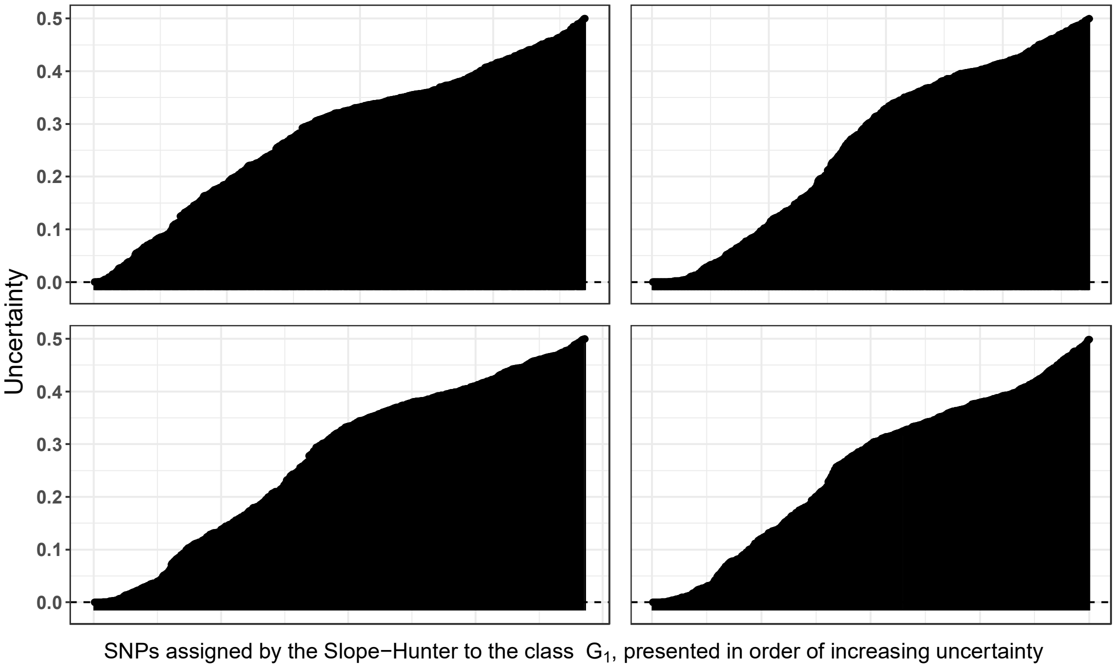


Figure S1: Uncertainty plot for SNPs identified by the Slope-Hunter method as the variants affecting incidence only from four simulations of scenario 6 in which 5% of SNPs have effects on incidence only and 10% on both incidence and prognosis, explaining 25% and 50% respectively of variation in incidence. 70% of the latter group were SNPs affecting both incidence and prognosis traits through the same exposure, explaining 35% of variation in incidence. The rest of SNPs in this group had uncorrelated effects on incidence and prognosis. The vertical black lines indicate misclassified SNPs
